# Morpho-physiological and gene expression responses of wheat by *Aegilops cylindrica* amphidiploids to salt stress

**DOI:** 10.1101/2020.06.07.139220

**Authors:** Razieh Kiani, Ahmad Arzani, S. A. M. Mirmohammady Maibody, Mehdi Rahimmalek, Khadijeh Razavi

**Affiliations:** Department of Agronomy and Plant Breeding, College of Agriculture, Isfahan University of Technology, 8415683111 Isfahan, Iran; Department of Horticulture, College of Agriculture, Isfahan University of Technology, 8415683111 Isfahan, Iran; National Institute of Genetic Engineering and Biotechnology (NIGEB), Tehran, Iran. P.O. Box: 14965/161, Tehran, Iran

**Keywords:** *HKT1;5*, *NHX1*, *SOS1*, *Ae. cylindrica*, amphidiploid, salt tolerance

## Abstract

*Aegilops cylindrica* Host is one of the most salt-tolerant species in the Triticeae tribe. Amphidiploid plants derived from hybridization of ‘Roshan’ × *Aegilops cylindrica* and ‘Chinese Spring’ × *Ae. cylindrica* genotypes contrasting in salt tolerance were assessed for their morpho-physiological responses and the expression patterns of three genes related to ion homeostasis under 250 mM NaCl. Results showed that salt stress caused significant declines in both their morphological and phenological traits. Moreover, salt stress reduced not only their chlorophyll content but also their root and shoot K contents and K/Na ratios, while it led to significant enhancements in the remaining traits. Similar to *Ae. cylindrica*, the amphidiploids subjected to salt stress exhibited significantly higher H_2_O_2_ levels, root and shoot K contents, and root and shoot K/Na ratios accompanied by lower root and shoot Na contents and MDA concentrations when compared with the same traits in the wheat parents. Quantitative Real-Time PCR showed significant differential expression patterns of the *HKT1;5, NHX1*, and *SOS1* genes between the amphidiploids and their parents. The transcript level of *HKT1;5* was found to be higher in the roots than in the shoots of both the amphidiploids and *Ae. cylindrica* while *NHX1* exhibited a higher expression in the shoot tissues. The consistency of these data provides compelling support for the hypothesis that active exclusion of Na from the roots and elevated vacuolar sequestration of Na in the leaves might explain the declining Na levels in the shoots and roots of both the amphidiploids and *Ae. cylindrica* relative to those measured in wheat parents. It is concluded that the hybridized amphiploids are potentially valuable resources for salt improvement in bread wheat through the backcrossing approach.

## Introduction

Soil salinity typically inhibits growth and impairs reproduction in plants through an initial osmotic-stress phase followed by ionic toxicity due to sodium (Na^+^) and chloride (Cl^−^) accumulation in the cell cytosol to lead ultimately to nutritional deprivation and oxidative stress (Arzani and Ashraf 2016; Munns and Tester 2008). Oxidative stress is caused by excessive amounts of such reactive oxygen species (ROS) as hydroxyl radical (OH.), superoxide radical (O_2_^-^), singlet oxygen (O_2_), and hydrogen peroxide (H_2_O_2_). The latter possibly plays a role in salt-stress by inducing K^+^ efflux from plant cells (Rubio et al. 2020). Briefly, activation of nonselective cation channels (NSCC) activates NADPH oxidase to increase the production of apoplastic hydrogen peroxide, whereby, NSCC-mediated K^+^ efflux is promptly promoted (Demidchik et al. 2007). This is further entrenched by the fact that halophytes employ such salt-tolerance mechanisms as restricting Na^+^ influx into the root, reduction of Na^+^ translocation, compartmentation of toxic ions into vacuoles, and sodium exclusion (Flowers and Colmer 2015; Maathuis et al. 2014; Arzani 2008).

The ability of photosynthetically active plant tissues to maintain intracellular Na and K homeostasis is one of their major salt stress tolerance mechanisms (Pan et al. 2020; Shabala et al. 2017; Shabala and Pottosin 2014; Wu et al. 2018). Impairment of the intracellular Na and K homeostasis, however, results in inhibited cell division, photosynthesis, growth, and development (Yan et al. 2013; Duan et al. 2015). K^+^/ Na^+^ ratio has been recognized as a critical physiological mechanism that contributes to the overall salt tolerance of plants (Rubio et al. 2020); in this mechanism Na^+^ does not accumulate to toxic levels in the leaves by discriminately excluding Na^+^ but accumulating K^+^ in the cells (Arabbeigi et al. 2014; Kiani et al. 2015; Shabala et al. 2013; Zhu et al. 2016). Recently, Arino-estrada et al. (2019) used the positron emission tomography (PET) to demonstrate the dynamic uptake and transport of Na in seedlings and to reveal the significant genetic diversity in terms of salt transport.

The high-affinity potassium transporters (HKTs) play a role in the long-distance transport of Na^+^ via phloem and xylem. In addition, the tonoplast-localized Na^+^/H^+^ antiporter (NHX1) (Apse et al. 2003), and the plasma membrane (PM)-localized salt-overly sensitive 1 (*SOS1*) Na^+^/H^+^ antiporter (Shi et al. 2002) excel at Na^+^ homeostasis. Munns et al. (2012) demonstrated the significant contribution of *Nax2* locus (*TmHKT1;5-A*) to reducing leaf Na^+^, that led to an improvement of 25% in durum wheat grain yield under saline soil conditions when compared with their near-isogenic lines lacking the *TmHKT1;5-A*.

A typical salt-excluding halophyte is the grass *Ae. cylindrica* with excellent salt tolerance. The intracellular and PM Na^+^/H^+^ exchangers (*NHX1* and *SOS1*, respectively), and the HKT-type protein (*HKT1;5*) are likely significant Na^+^ transporters contributing to salt tolerance in this species (Arabbeigi et al. 2018). Motivated by our previous works of screening the hyper-salt tolerant genotypes of *Ae. cylindrica*, (Kiani et al. 2015), hybridizing wheat and a hyper salt-tolerant *Ae. cylindrica* accompanied by cytological and molecular assessments of the synthesized amphidiploid plants (Kiani et al. unpublished data), and that of Arabbeigi et al. (2018) who tested and identified the expression patterns of some of the most important genes involved in Na^+^ homeostasis in *Ae. cylindrica*, the current study was implemented to investigate the physiological responses and expression patterns of the genes *HKT1;5, NHX1* and *SOS1* in two wheat cultivars, one genotype of *Ae. cylindrica*, and their interspecific hybrid derivatives when exposed to 250 mM NaCl.

## Materials and methods

### Plant materials

Wheat × *Ae. cylindrica* amphidiploid plants were developed through hybridizing wheat (*T. aestivum* L.) cultivars (‘Chinese Spring’ and ‘Roshan’) and *Ae. cylindrica*. For this purpose, a hyper-salt-tolerant genotype of *Ae. cylindrica* (USL26) originated from the shores of the Urumia Salt Lake in Northwest Iran (Arabbeigi et al. 2014; Kiani et al. 2015) was used as the male parent. Before and after pollination (24 hr each), the uppermost internodes were injected with a mixed solution (1:1) of 25 mg L^−1^ gibberellic acid (GA_3_) and 10 mg L^−1^ 2.4-dichlorophenoxyacetic acid (2,4-D). Fourteen days after pollination, the immature embryos were cultured in the Murashige and Skoog (MS) basal medium (Sigma-Aldrich M9274) containing 0.8% agar and 2% sucrose. Subsequently, the colchicine treatment (Nemeth et al. 2015; Arzani and Darvey 2001) was carried out at the 4-5 tiller stage. The roots were then placed in a 0.1% (w/v) solution of colchicine supplemented with 2% DMSO (dimethyl sulfoxide) and 2 drops of Tween-20 per 100 mL of the solution at the 2-4 tiller stage for 5 h at 20° C. The roots were ultimately washed with tap water for 1 h before being grown under greenhouse conditions (Arzani and Darvey 2001).

### Field experiment

A field experiment was carried out during 2017–2018 at the research farm of Isfahan University of Technology, located at Lavark (32–300N, 51–200E; 1,630 m asl), Iran, with a mean annual temperature and precipitation of 15° C and 140 mm. Prior to sowing, surface soil samples from the experimental site were taken from a depth of 0–30 cm, air-dried, and sieved through a 2-mm screen in order to determine their physico-chemical characteristics through analysis in the laboratory according to standard methods. The top 60-cm layer was found to be a clay loam soil (pH 7.5) with a bulk density of 1.48 g/cm3 and an average EC of 4 dS m^−1^ at the sowing date. At the harvesting date, however, the average soil EC of the salt-treated experimental plots was measured at 20.4 dS m^−1^. A split-plot design with three replications incorporating a randomized complete block arrangement was used. Salt treatments (normal/control and 250 MM NaCl) were assigned to the main plots and the genotypes to the sub-plots. Each plot contained two rows 2-m long spaced 30 cm from each other with plants spaced 10 cm from each other within the rows. Irrigation was applied via a drip system throughout the period from sowing to the late grain-filling stage. Salt treatment was initiated at the 4-tiller stage using initially an irrigation water containing diluted NaCl (125 mM) but continued with raising NaCl concentration to 250 mM.

### Morphological traits

Agronomic traits including number of tillers, plant height, number of spikelets per spike (spikelet/spike), spike length, number of spikes per plant (spike/plant), flag leaf width (FLW), flag leaf length (FLL), flag leaf area (FLA), fresh flag leaf weight (FFLW), dry flag leaf weight (DFLW), plant fresh weight (FW), and plant dry weight (DW) were evaluated under the two control and salt stress conditions. The number of productive tillers per plant was determined prior to harvesting. Plant height from the ground level to the spike tip, excluding the awns, was recorded at maturity. FLL, FLW, and FLA were recorded using a leaf area instrument (Fanavaran Alborz Andisheh Co. Iran). FFLW and FW were determined using ten randomly selected plants. Finally, DFLW and DW were recorded using fresh samples dried in an oven at 70° C for 48 h.

### Phenological traits

Days to heading (DH) defined as days from sowing to the emergence of 75% of the heads (spikes) in a plot, days to pollination (DP) representing the number of days from sowing to the emergence of anthers in 75% of the plants in a plot, and days to maturity (DM) denoting the number of days from planting to physiological maturity when the peduncles turned yellow were recorded under both control and salt-stress conditions.

### Physiological traits

#### Enzymatic antioxidants

For enzyme extraction, 0.1 g of leaf sample was ground into a fine powder using liquid nitrogen and transferred into the extraction buffer containing 50 mM sodium phosphate (pH 7.8), 1% polyvinylpolypyrrolidone (PVP), 0.2% Triton X-100, 2 mM EDTA, 2 mM dithiothreitol (DTT), and 50 mM Tris–HCl. The homogenate was subsequently centrifuged at 12,000 g for 30 min before the supernatant was decanted and assayed for enzyme activity.

Ascorbate peroxidase (APX) activity was determined as the decrease in absorbance at 290 nm for 2 min (Nakano and Asada 1981). Catalase (CAT) activity, expressed as mmol H_2_O_2_ min^−1^ mg^−1^ of protein, was evaluated by measuring the rate of decrease in H_2_O_2_ absorbance at 240 nm as previously described (Aebi, 1984). The activity of guaiacol peroxidase (GPX) was assessed as the increase of absorbance at 470 nm for 2 min (Herzog and Fahimi 1973). Finally, protein content was determined by using bovine serum albumin as the standard (Bradford 1976).

#### Lipid peroxidation (MDA)

The thiobarbituric acid (TCA) test was used to measure malondialdehyde (MDA) as the lipid peroxidation product in leaf homogenates (Taulavuori et al. 2001). Briefly, 0.30 g of fresh leaf sample was homogenized in 5 ml of 0.1% TCA. The homogenate was then centrifuged at 12,000 g for 10 min at 4° C. Four ml of 20% TCA containing 0.5% thiobarbituric acid (TBA) was mixed in 1 ml of the supernatant, heated for 30 min at 95° C in a water bath, and immediately cooled on ice. After centrifugation at 10,000 g for 10 min, the absorbance of the extract was read at 532 nm and the values were corrected for non-specific turbidity by subtracting from absorbance at 600 nm. An extinction coefficient of 155 mM^−1^ cm^−1^ was used to calculate the MDA content using the following Equation:

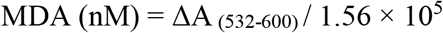

#### Hydrogen peroxide (H_2_O_2_)

Hydrogen peroxide (H_2_O_2_) was determined following the procedure proposed in Velikova et al. (2000). For this purpose, fresh leaf tissue (0.30 g) was homogenized in 5 ml TCA. The homogenate was subsequently centrifuged at 12,000 g for 10 min at 4° C and the supernatant was neutralized with 5 M K_2_CO_3_ (pH =5.6) in the presence of 100 μL of 0.3 M phosphate buffer (pH=5.6). The reaction was allowed to continue for 1 h in the dark; absorbance was ultimately measured at 390 nm. H_2_O_2_ concentration was calculated using the standard curve based on known concentrations of H_2_O_2_.

#### Carotenoid and chlorophyll content

The chlorophyll (a+b) and carotenoid (x+ c) contents of leaf tissues (0.33 g) in 80% acetone extract were determined spectrophotometrically using the redetermined extinction coefficients and equations introduced by Lichtenthaler and Buschman (2001). Briefly, about 0.5 g of fresh flag leaf was cut from mid-section, ground and mixed in 5 mL of 80% acetone, and stored in the dark for 5 min. The solution was then filtered and the absorbance of the filtrate was read at 645, 663, and 470 nm using 80% acetone as the blank. The quantities of chlorophyll a, chlorophyll b, total chlorophyll, and carotenoids were calculated and expressed in mg g^−1^ FW using the following equations:

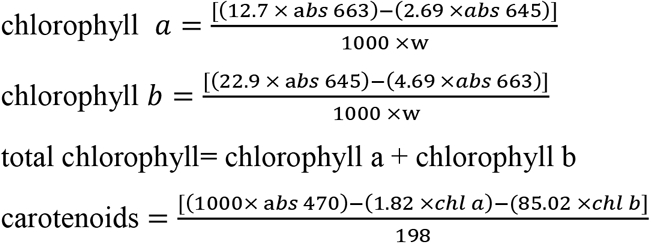

#### Leaf and root Na and K contents

Root and shoot samples were dried for 48 h at 80° C to constant weights. To determine leaf and root Na and K contents, relevant samples (0.2 g dry weight) were incinerated in a muffle furnace at 550° C for 4 h. The ash thus obtained was then dissolved in 10 ml of 2 N HCl and its volume was made to 100 ml. Using a standard curve, the mineral contents were determined by flame photometry (Jenway PFP7, UK) (Houshmand et al. 2005) and the K/Na ratio was accordingly calculated.

#### Gene expression analysis

The sown seeds were allowed to germinate for 7 days at 4° C before they were transferred into pots 30 cm in height and 20 cm in diameter, which had been filled with a mixture of soil and sand (coarse crushed) at a ratio of 3:1 and placed in a greenhouse with average day/night temperatures of 25 ± 3/17 ± 3 °C, respectively; a relative humidity of 60–65%; and a photoperiod of 12 h with a photosynthetic photon flux density (PPFD) of about 480 μmol m^−2^ s^−1^. The seedlings were initially watered with tap water, which was continued throughout the experiment only for the control pots (EC = 1.5 dS m^−1^). For the salt treatment pots, however, salt stress was initiated by gradually adding 250 mM NaCl upon the emergence of the second leaf. A three-factor factorial experiment was conducted to assess the effects of NaCl treatments (0 and 250 mM NaCl at 24 h and 7 days for the gene expression assay) on two tissue types (root and shoot) of five genotypes with four replicates (two for each of the biological and technical) of 10 plants. Roots and shoots from each experimental unit were sampled, snap-frozen in liquid nitrogen, and immediately stored at –80° C until use.

#### RNA extraction and cDNA synthesis

The plant tissue sample (100 mg) was ground into a fine powder using liquid nitrogen. Total RNA was isolated using the CTAB method (Murray and Thompson 1980) with minor modifications. Briefly, the powder was homogenized with the extraction buffer (2% CTAB, 20 mM EDTA pH 8.0, 2% PVP-40, 0.1 M Tris-HCl pH 8.0, 1.4 M NaCl, and 10% of β-mercaptoethanol added immediately prior to use) and the tube was incubated at 65° C for 10 min. Eight milliliters of chloroform was then added to the tube before its vigorous inversion and centrifugation at 10,000 g for 10 min at 4° C. The supernatant was transferred to a new tube, mixed with an equal volume of chloroform:phenol (1:1), and centrifuged at 10000×g for 10 min at 4° C. An equal volume of chloroform:isoamyl alcohol (24:1 v/v) was added to the supernatant obtained from centrifugation, which was then mixed and centrifuged at 10,000 g for 10 min at 4° C. Subsequently, the supernatant was mixed with LiCl (8M ¼ vol concentration) in a new tube and incubated at –20° C for 4 hours. The precipitated RNA was pelleted by centrifugation at 12,000 g at 4° C for 20 min, the pellets were subsequently washed twice with ice-cold 70% ethanol, dried, and finally suspended in RNase-free water.

DNase treatment of total RNA was carried out by Sina kit, Iran, according to the manufacturer’s instructions. The purity of the total RNA was quantified using an ND-1000 NanoDrop (Germany). Synthesis of cDNA from l µg RNA was carried out using Suprime - Script RT premix (GeNet Bio, Korea) according to the manufacturer’s instructions.

#### Quantitative Real-Time RT-PCR

Using the gene-specific primers of the *HKT1;5, NHX1* and *SOS1* genes (namely, 5’-TCCTCCTTG\GACTTGATGTAG-3’/5’-CCATAGACCTTGACCTGAT-3’; 5’-GAATGCCACTCAGATCCAGC-3’/5’-GCTGCTGGGTGGCTTAGTGC-3’; and 5’-CAAGGATTTTGCTCCCTCAG-3’/5’-CTTCTGGTTCGCTTCCACAT-3’, respectively), the expression profiles of the above genes were analyzed by conducting qRT-PCR in 20 µl of a reaction mixture containing 1 µl of diluted cDNA, 5 µl of 2x Fast SYBR Green PCR Master Mix (Applied Biosystems), 0.2 µl of 1 µM each of the reverse and forward primers, and 13.6 µl of RNAase free water. Amplification was performed on a StepOnePlusTM Real-Time PCR system (Applied Biosystems) according to the following cycling protocol: 94° C for 4 min, 42 cycles of 95° C for 30 s, 60° C for 35 s, 72° C for 1.5 min, and 72° C for 10 min. A housekeeping gene, β-actin (ACTB), was used as the reference to normalize the expressions of the genes studied. The data of real-time RT-PCR were analyzed using the 2^-ΔΔCT^ method (Livak and Schmittgen 2001). The relative expression levels of the target mRNA in both roots and shoots were calculated using the ΔΔCT method and expressed relative to the values in the sham (normal tissues) after normalization.

### Statistical analysis

The data obtained were tested for homogeneity and normality of variance before being subjected to analysis of variance (ANOVA) using SAS version 9.3 (SAS Institute 2011) in which genotype and treatment were considered as fixed effects and replication as a random effect. The Fisher’s LSD test (LSD_0.05_) was used to detect significant differences among the means of the treatments. Correlation and linear regression analyses were performed to delineate the relationships among the traits using SAS PROC CORR and SAS PROC REG, respectively. Principal component analysis (PCA) was carried out on the data on physiological responses to salt stress (250 mM NaCl) obtained for the amphidiploids and their parents; the Kaiser’s rule of Eigenvalue exceeded 1.0 using PROC FACTOR (method = prin) in SAS 9.3. The first two PCs (PC1 and PC2) were plotted to construct the genotype-by-trait biplot using Stat Graphics.

## Results

### Morphological traits

The results of ANOVA showed that salt stress had significant effects on all the traits except the number of spikelets per spike, spike length, number of spikes per plant, FLL, and FLW (Table 1). The overall means of the number of tillers and plant height were observed to reduce by 26.60% and 17.86%, respectively, while the overall means of FW and DW decreased by 49.54% and 65.61 %, respectively, in 250 mM NaCl treatment (Table 2). Significant interaction effects of salt stress by genotype were only detected for the number of tillers, FW, and DW traits (Table 1).

**Table 1:** Results of analysis of variance for the traits studied under the control and salt-stress (250 mM NaCl) treatments

| Trait | Mean square |  |  |  |  |
| --- | --- | --- | --- | --- | --- |
|  | Salt stress (S) | Genotype (G) | G × S | Residual | CV % |
| **Morphological traits** |  |  |  |  |  |
| Number of tillers | 267.94** | 395.04** | 67.14** | 7.71 | 14.25 |
| Plant height | 1378.75** | 1371.46** | 49.72<sup>ns</sup> | 41.65 | 9.33 |
| Spikelet/spike | 140.83<sup>ns</sup> | 478.11** | 164.26<sup>ns</sup> | 53.41 | 20.55 |
| Spike length | 0.08<sup>ns</sup> | 11.97** | 0.99<sup>ns</sup> | 0.28 | 5.49 |
| Spike/plant | 25.17<sup>ns</sup> | 169.65** | 1.23<sup>ns</sup> | 10.34 | 27.79 |
| FLL | 2.61<sup>ns</sup> | 8.02* | 5.90<sup>ns</sup> | 2.42 | 9.88 |
| FLW | 0.002<sup>ns</sup> | 1.41* | 0.03<sup>ns</sup> | 0.02 | 12.62 |
| FLA | 46.33* | 66.05** | 19.63<sup>ns</sup> | 6.63 | 19.15 |
| FFLW | 0.01* | 0.05** | 0.02<sup>ns</sup> | 0.003 | 16.04 |
| DFLW | 0.006* | 0.008** | 0.002<sup>ns</sup> | 0.001 | 22.52 |
| FW | 0.01* | 0.01* | 0.01** | 0.002 | 13.64 |
| DW | 0.01* | 0.05** | 0.003** | 0.001 | 16.60 |
| **Phenological traits** |  |  |  |  |  |
| DH | 93.63** | 498.22** | 3.38<sup>ns</sup> | 7.87 | 1.76 |
| DP | 177.63** | 195.53** | 14.46<sup>ns</sup> | 13.53 | 2.19 |
| DM | 589.63** | 207.03** | 21.47* | 6.23 | 1.38 |
| **Physiological traits** |  |  |  |  |  |
| Chl<sub>a</sub> | 0.001** | 0.003* | 0.0002<sup>ns</sup> | 0.0002 | 15.86 |
| Chl<sub>b</sub> | 0.003** | 0.001** | 0.0006<sup>ns</sup> | 0.0003 | 38.06 |
| Tchl | 0.009** | 0.002** | 0.0005<sup>ns</sup> | 0.0009 | 21.08 |
| Car | 15.07** | 4.86** | 1.77<sup>ns</sup> | 1.40 | 11.06 |
| CAT | 0.074<sup>ns</sup> | 0.22* | 0.30<sup>ns</sup> | 0.098 | 27.29 |
| APX | 0.02<sup>ns</sup> | 0.02<sup>ns</sup> | 0.01<sup>ns</sup> | 0.02 | 51.62 |
| GPX | 1.01* | 0.52* | 0.62<sup>ns</sup> | 0.29 | 43.18 |
| H<sub>2</sub>O<sub>2</sub> | 0.45* | 0.43* | 0.10<sup>ns</sup> | 0.13 | 26.14 |
| MDA | 0.194** | 0.021** | 0.01<sup>ns</sup> | 0.008 | 33.65 |
| Leaf Na | 0.28** | 0.04** | 0.002<sup>ns</sup> | 0.001 | 3.21 |
| Leaf K | 12.86** | 0.46** | 0.42** | 0.06 | 9.678 |
| Leaf K/Na | 7.65** | 0.34** | 0.02<sup>ns</sup> | 0.03 | 13.24 |
| Root Na | 11.08** | 0.11* | 0.06<sup>ns</sup> | 0.03 | 8.76 |
| Root K | 4.87** | 0.09** | 0.54** | 0.02 | 5.39 |
| Root K/Na | 15.17** | 0.16* | 0.18* | 0.04 | 13.60 |

Number of spikelets per spike (spikelet/spike), number of spikes per plant (spike/plant), flag leaf length (FLL), flag leaf width (FLW), flag leaf area (FLA), fresh flag leaf weight (FFLW), dry flag leaf weight (DFLW), plant fresh weight (FW), plant dry weight (DW), days to heading (DH), days to pollination (DP), days to maturity (DM), chlorophyll a (Chl<sub>a</sub>), chlorophyll b (Chl<sub>b</sub>), total chlorophyll (Tchl), carotenoid (Car), catalase (CAT), ascorbate peroxidase (APX), guaiacol peroxidases (GPX), MDA and H<sub>2</sub>O<sub>2</sub> content, root and shoot Na and K contents and K/Na ratios.
<sup>ns</sup> non-significant, \* $p < 0.05$ and \*\* $p < 0.01$

**Table 2:**
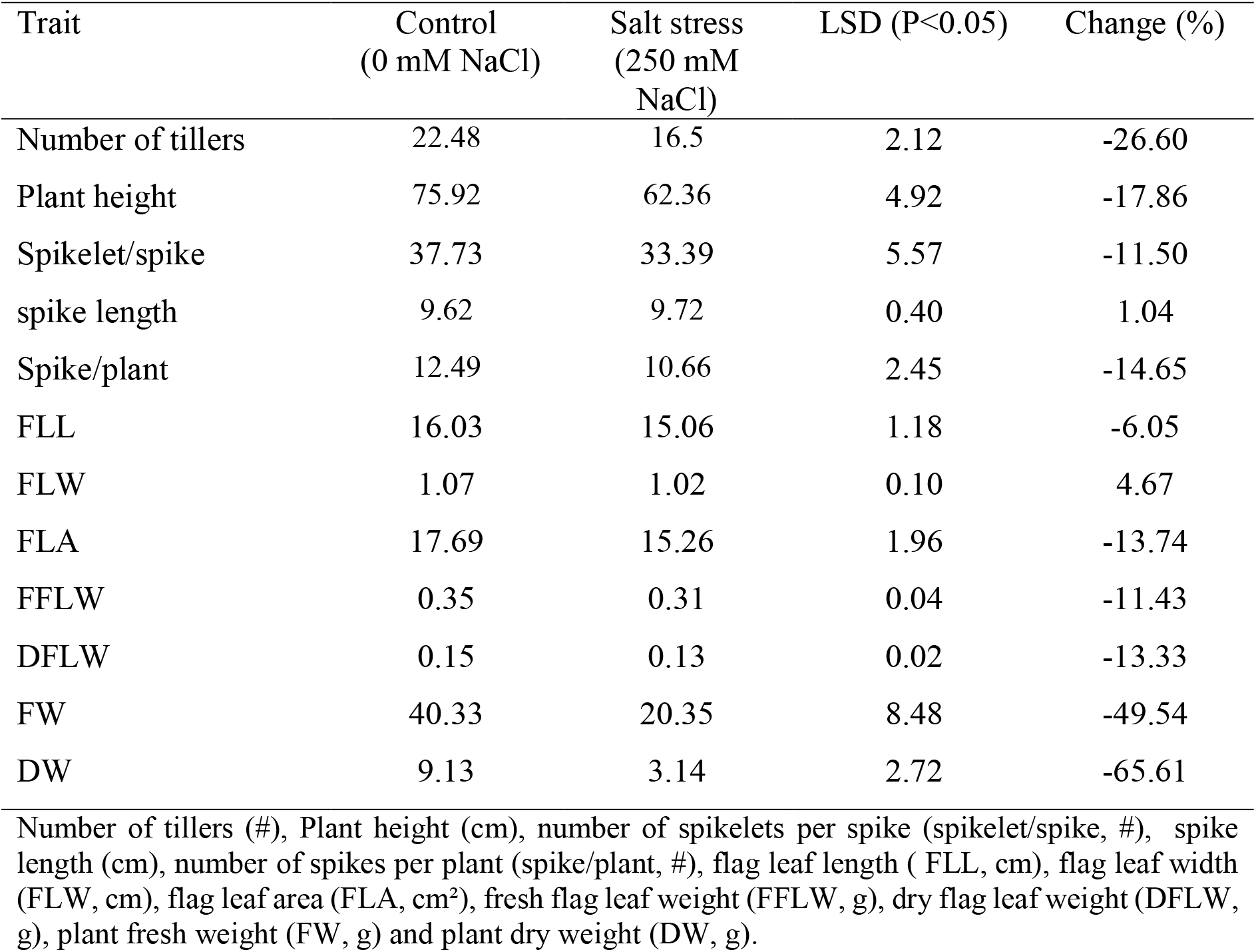
Mean comparisons of morphological traits under the control and salt-stress (250 mM NaCl) conditions

| Trait | Control<br>(0 mM NaCl) | Salt stress<br>(250 mM<br>NaCl) | LSD (P<0.05) | Change (%) |
| --- | --- | --- | --- | --- |
| Number of tillers | 22.48 | 16.5 | 2.12 | -26.60 |
| Plant height | 75.92 | 62.36 | 4.92 | -17.86 |
| Spikelet/spike | 37.73 | 33.39 | 5.57 | -11.50 |
| spike length | 9.62 | 9.72 | 0.40 | 1.04 |
| Spike/plant | 12.49 | 10.66 | 2.45 | -14.65 |
| FLL | 16.03 | 15.06 | 1.18 | -6.05 |
| FLW | 1.07 | 1.02 | 0.10 | 4.67 |
| FLA | 17.69 | 15.26 | 1.96 | -13.74 |
| FFLW | 0.35 | 0.31 | 0.04 | -11.43 |
| DFLW | 0.15 | 0.13 | 0.02 | -13.33 |
| FW | 40.33 | 20.35 | 8.48 | -49.54 |
| DW | 9.13 | 3.14 | 2.72 | -65.61 |

Number of tillers (#), Plant height (cm), number of spikelets per spike (spikelet/spike, #), spike length (cm), number of spikes per plant (spike/plant, #), flag leaf length ( FLL, cm), flag leaf width (FLW, cm), flag leaf area (FLA, cm<sup>2</sup>), fresh flag leaf weight (FFLW, g), dry flag leaf weight (DFLW, g), plant fresh weight (FW, g) and plant dry weight (DW, g).

Salt stress was found to cause a significant reduction in growth and morphological traits (Table 3), whilst the amphidiploids derived from both ‘Chinese Spring’ × *Ae. cylindrica* and ‘Roshan’ × *Ae. cylindrica* crosses were less affected compared to the wheat cultivars (i.e., female parents) but more affected than their wild male parent. The F_1_ amphidiploid plants derived from the ‘Chinese Spring’ × *Ae. cylindrica* cross recorded average numbers of tillers equal to 33 and 24.06 tillers per plant under control and salt stress conditions, respectively, whereas the ‘Chinese Spring’ recorded a mean number of 12.61 tillers per plant in the control and 9.06 in the salt stress treatment. Although the highest reduction in plant height was observed in ‘Chinese spring’ (16.18), both *Ae. cylindrica* and ‘Chinese Spring’ × *Ae. Cylindrica* hybrid plants were affected by salt stress as evidenced by reductions of 13.19% in *Ae. cylindrica* and 14.02% in ‘Chinese spring’ × *Ae. cylindrica* amphidiploids (Table 3). Drastic reductions of 50.91% and 42.71% due to salt stress were observed in the number of spikes per plant in the female parents (wheat cultivars), while the number of spikelets per spike and spike length were comparatively less affected (Table 3). On the other hand, salt stress caused only a minor and insignificant reduction in these traits in either *Ae. cylindrica* or the amphidiploid plants. Likewise, the highest reduction in flag leaf area (27.38%) due to salt stress was observed in ‘Chinese Spring’ (Table 3). Comparisons based on DFLW indicated no significant changes in the case of *Ae. cylindrica* and ‘Chinese Spring’ × *Ae. cylindrica* hybrid plants as a result of salt stress, while considerable reductions (29.41 % and 26.32 %, respectively) were observed in ‘Chinese Spring’ and ‘Roshan’ as the female parents (Table 3). Although all the genotypes exhibited significantly decreased FW and DW values due to salt stress, lower reductions in these traits were observed in ‘Roshan’ × *Ae. cylindrica*, ‘Chinese Spring’ ×*Ae. cylindrica* amphidiploids, and their male parent (*Ae. cylindrica*) under salt stress (Table 3).

**Table 3:** Mean comparisons of morphological traits of the parent plants (‘Roshan’, ‘Chinese spring’, *Ae. cylindrica*) and their interspecific F_1_ hybrids (amphidiploids) under control and salt-stress (250 mM NaCl) conditions

| Trait | Genotype |  |  |  |  |  |  |  |  |
| --- | --- | --- | --- | --- | --- | --- | --- | --- | --- |
|  | ‘Roshan’ |  |  | ‘Chinese spring’ |  |  | *Ae. cylindrica* |  |  |
|  | Control | Stress | Change (%) | Control | Stress | Change (%) | Control | Stress | Change (%) |
| Number of tillers | 14.67 | 13.84 | -5.66 | 12.61 | 9.06 | -28.15 | 35.00 | 29.33 | -16.20 |
| Plant height | 85.09 | 72.23 | -15.11 | 84.26 | 70.63 | -16.18 | 48.00 | 41.67 | -13.19 |
| Spikelet/spike | 44.38 | 41.33 | -6.87 | 37.40 | 35.80 | -4.28 | 22.67 | 20.33 | -10.32 |
| Spike length | 10.87 | 10.63 | -2.21 | 7.97 | 7.50 | -5.90 | 10.78 | 10.28 | -4.64 |
| Spike/plant | 11.59 | 5.69 | -50.91 | 6.79 | 3.89 | -42.71 | 21.67 | 16.33 | -24.64 |
| FLL | 14.93 | 14.75 | -1.21 | 16.70 | 13.02 | -22.04 | 17.70 | 16.98 | -4.07 |
| FLW | 1.58 | 1.37 | -13.29 | 1.27 | 1.23 | -3.15 | 0.59 | 0.51 | -13.56 |
| FLA | 17.95 | 14.70 | -18.11 | 12.27 | 8.91 | -27.38 | 10.49 | 8.66 | -17.45 |
| FFLW | 0.42 | 0.35 | -16.67 | 0.36 | 0.32 | -11.11 | 0.21 | 0.18 | -14.29 |
| DFLW | 0.19 | 0.14 | -26.32 | 0.17 | 0.13 | -29.41 | 0.10 | 0.09 | -10.00 |
| FW | 40.42 | 22.34 | -44.73 | 40.36 | 20.32 | -49.65 | 28.20 | 17.17 | -39.11 |
| DW | 12.17 | 4.13 | -66.06 | 10.17 | 3.14 | -69.12 | 7.08 | 3.46 | -51.13 |

| Trait | ‘Roshan’ × *Ae. cylindrica*<br>amphidiploid |  |  | ‘Chinese spring’ × *Ae. cylindrica*<br>amphidiploid |  |  | LSD (P<0.05) |  |
| --- | --- | --- | --- | --- | --- | --- | --- | --- |
|  | Control | Stress | Change (%) | Control | Stress | Change (%) | Control | Stress |
| Number of tillers | 18.24 | 16.21 | -11.13 | 33.00 | 24.06 | -27.09 | 4.89 | 5.87 |
| Plant height | 78.91 | 67.67 | -14.24 | 69.33 | 59.61 | -14.02 | 11.11 | 12.44 |
| Spikelet/spike | 41.33 | 38.50 | -6.85 | 30.87 | 21.00 | -31.97 | 21.15 | 3.37 |
| Spike length | 10.13 | 9.54 | -5.82 | 9.93 | 9.16 | -7.75 | 1.32 | 0.60 |
| Spike/plant | 11.42 | 8.22 | -28.02 | 16.27 | 12.14 | -25.38 | 4.08 | 6.74 |
| FLL | 15.19 | 14.98 | -1.38 | 16.55 | 16.48 | -0.42 | 3.24 | 1.77 |
| FLW | 1.48 | 1.33 | -10.14 | 0.64 | 0.62 | -3.13 | 0.26 | 0.26 |
| FLA | 15.47 | 14.49 | -6.33 | 17.27 | 14.28 | -17.31 | 5.38 | 4.31 |
| FFLW | 0.36 | 0.31 | -13.89 | 0.35 | 0.31 | -11.43 | 0.11 | 0.08 |
| DFLW | 0.15 | 0.13 | -13.33 | 0.19 | 0.17 | -10.53 | 0.08 | 0.04 |
| FW | 30.36 | 19.29 | -36.42 | 30.38 | 16.21 | -46.64 | 3.16 | 2.19 |
| DW | 9.17 | 4.11 | -55.18 | 8.19 | 3.31 | -59.58 | 0.07 | 0.07 |

Number of tillers (#), Plant height (cm), Number of spikelets per spike (spikelet/spike, #), spike length (cm), number of spikes per plant (spike/plant, #), flag leaf length (FLL, cm), flag leaf width (FLW, cm), flag leaf area (FLA, cm<sup>2</sup>), fresh flag leaf weight (FFLW, g), dry flag leaf weight (DFLW, g), plant fresh weight (FW, g) and plant dry weight (DW, g).

### Phenological traits

The amphidiploid plants derived from the ‘Roshan’ × *Ae. cylindrica* and ‘Chinese Spring’ × *Ae. cylindrica* crosses as well as their parents exhibited significant changes in phenological traits (DH, DP, and DM) when subjected to 250 mM NaCl stress compared to those grown under the control conditions (Table 1). The amphidiploid plants derived from ‘Roshan’ × *Ae. cylindrica* and ‘Chinese Spring’ × *Ae. cylindrica* crosses showed significant declines in DH due to salt stress, with mean values of 161.33 and 157.80 days under the control and salt stress conditions, respectively (Fig. 1). Moreover, the average values 170.13 and 185.73 days obtained for DP and DM under the control conditions declined to 165.27 and 176.87 days, respectively, under salt stress (Fig. 1).

**Fig 1:**
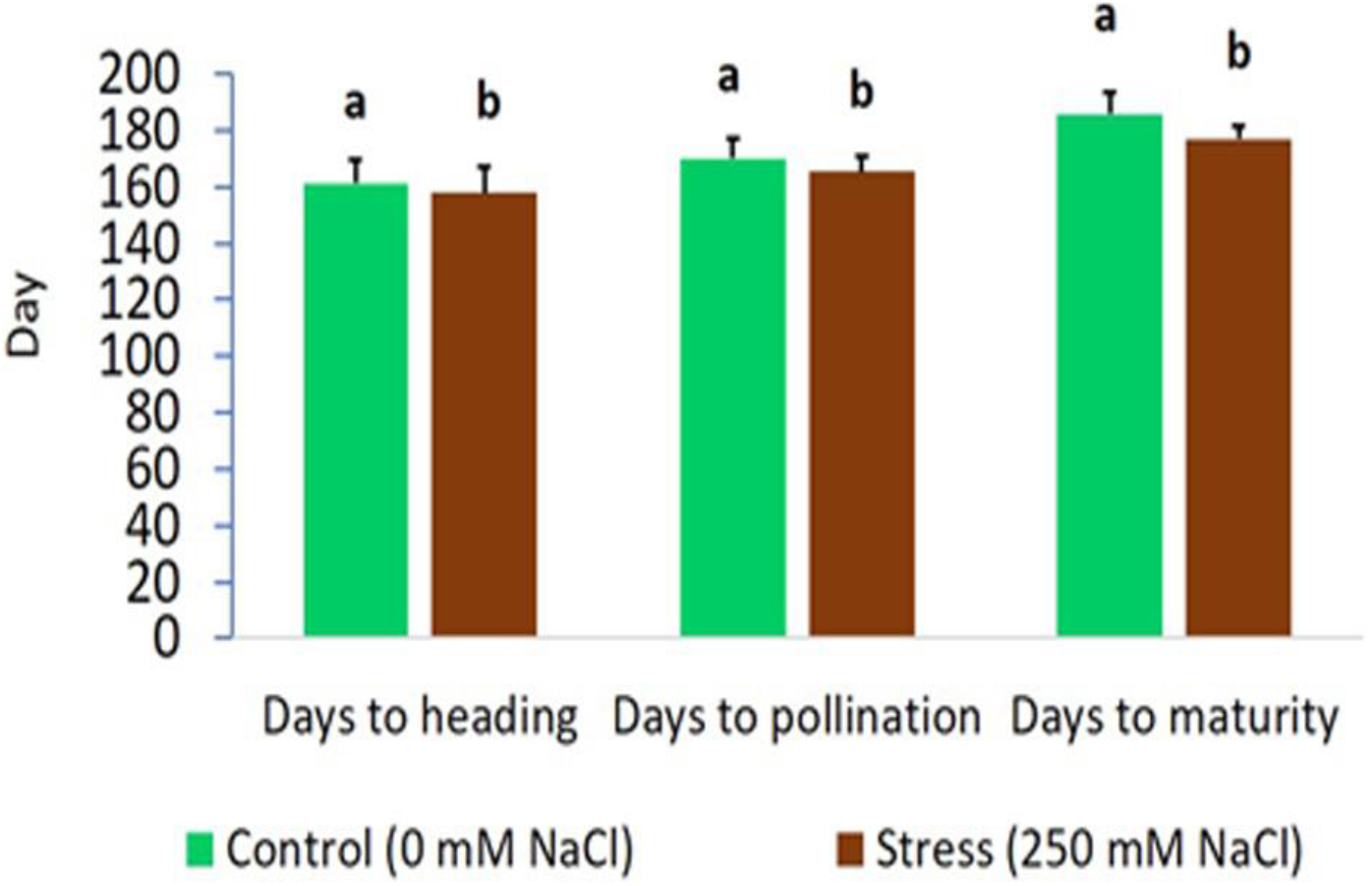
Effects of salt stress on days to heading, days to pollination, and days to maturity in parents (‘Roshan’, ‘Chinese spring’, *Ae. cylindrica*) and their F_1_ hybrid plants. Bars represent means ± SE and bars with the same letter do not significantly differ at *P*<0.05.

Compared to the common wheat (*T. aestivum* L.) cultivars (i.e., ‘Roshan’ and ‘Chinese Spring’), *Ae. cylindrica* exhibited lower DH, DP, and DM values in both control and salt stress conditions (Fig. 2). However, the interspecific hybrid plants obtained from the ‘Roshan’ × *Ae. cylindrica* and ‘Chinese Spring’ × *Ae. cylindrica* crosses exhibited intermediate phenology (Fig. 2).

**Fig 2:**
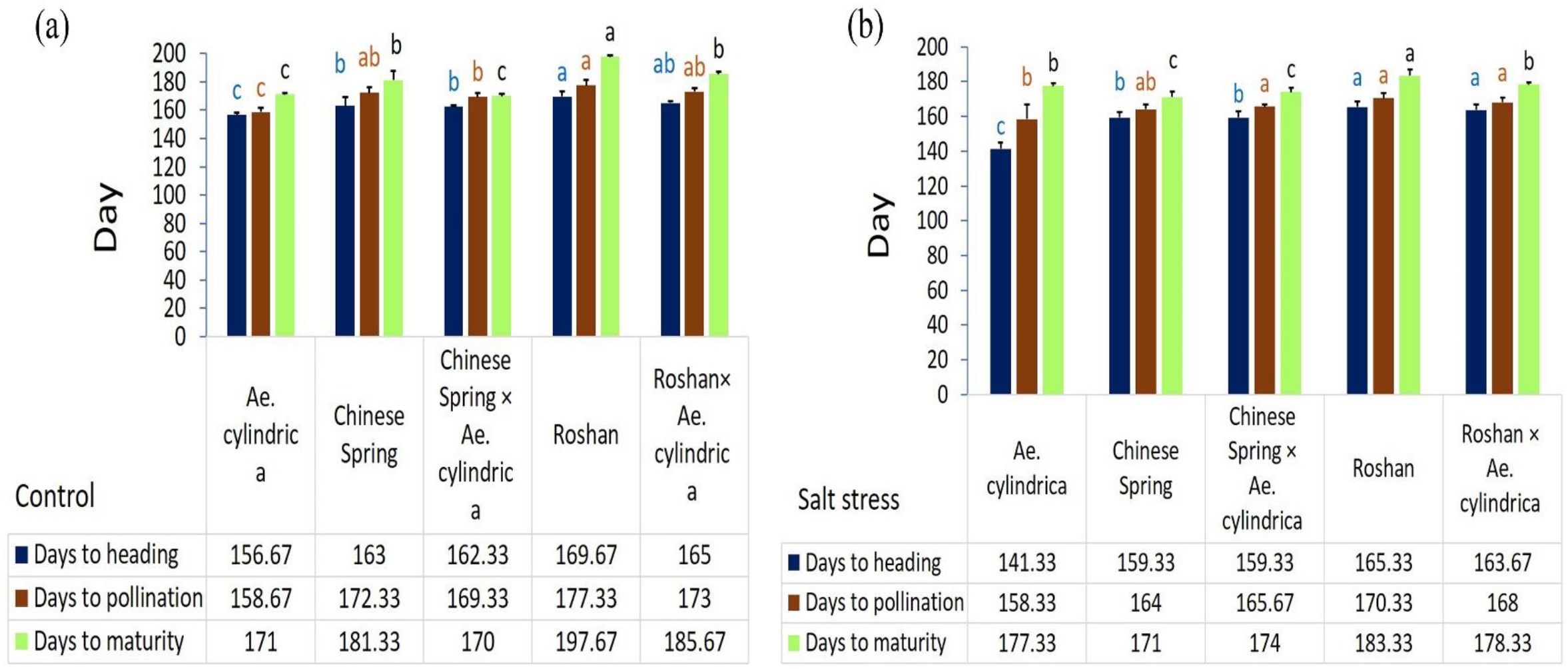
Mean comparison of phenological traits of parents (‘Roshan’, ‘Chinese spring’, *Ae. cylindrica*) and their F_1_ hybrid plants (amphidiploids) under (A) control and (B) salt stress (250 mM NaCl) conditions. Bars represent means ± SE and bars with the same letter do not significantly differ at *p*<0.05.

### Physiological traits

Results of the analysis of variance showed that all the physiological traits except CAT and APX activities were significantly affected (p<0.01) by salt stress (Table 1). The genotypes differed significantly with respect to the studied physiological traits except for APX activity (Table 1). Salt stress led to significantly increased GPX activity, Chl_a_, Car, MDA, and H_2_O_2_ content (Fig. 3), while the other two antioxidant enzymes (namely, APX and CAT) showed no significant effects of salt stress (Fig. 3a). Compared to the control conditions, salt stress led to increases of 26.82% and 7.09% in the overall means of Chl_a_ and Car, respectively (Fig. 3b and c). The highest increases in Chl_a_ (27.10 %), Tchl (3.35 %), and Car (10.52%) due to salt stress were recorded for *Ae. cylindrica* (Table 4). In addition, *Ae. cylindrica* and ‘Chinese spring’ exhibited the lowest and highest reductions in Chl_b_ content (31.94 % and 42.62%, respectively) under salt stress when compared with recorded values in the control treatment.

**Table 4:**
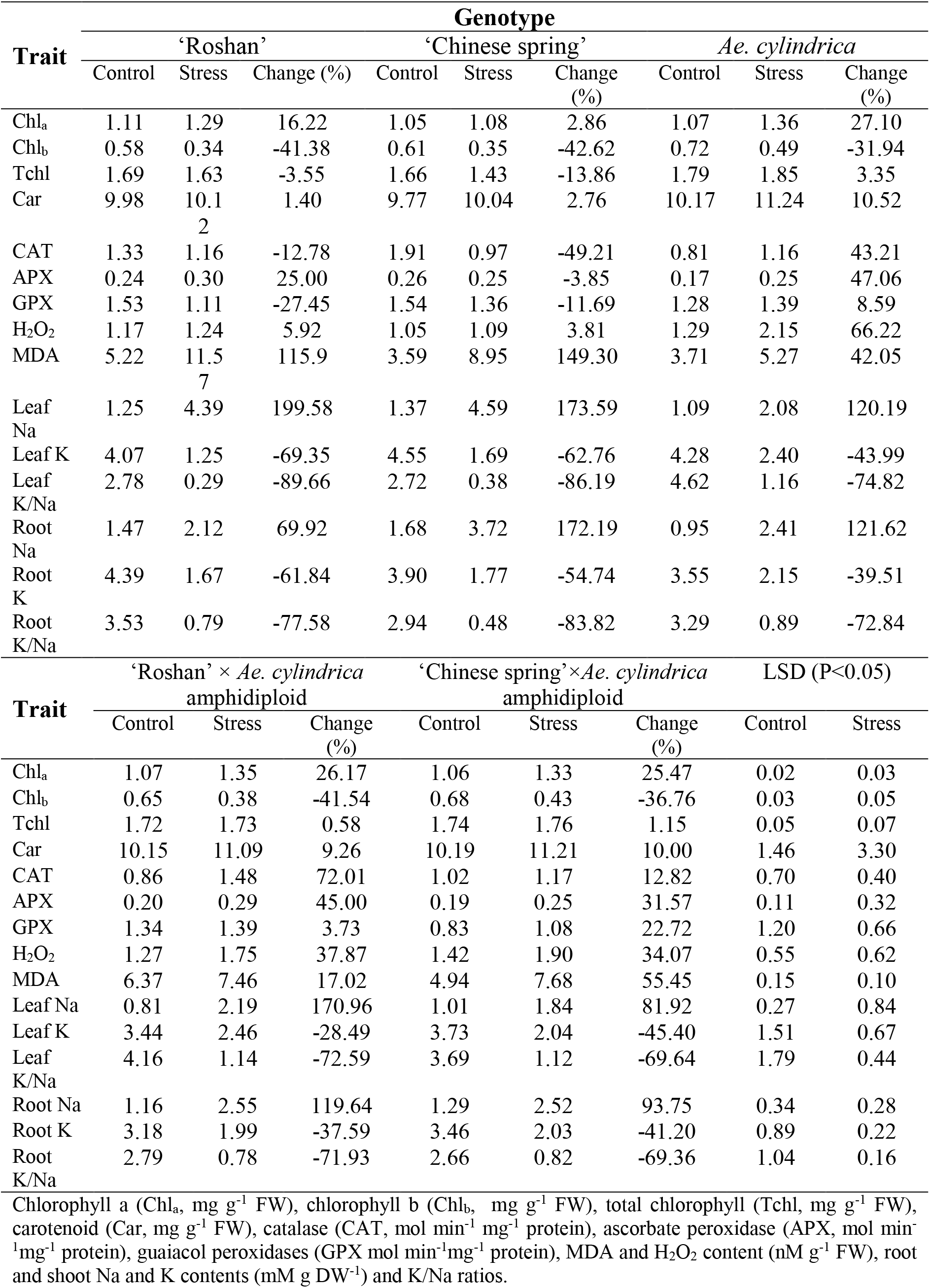
Mean comparisons of physiological traits of the parent plants (‘Roshan’, ‘Chinese spring’, *Ae. cylindrica*) and their F_1_ hybrid plants (amphidiploids) under the control and salt-stress (250 mM NaCl) treatments

**Fig 3:**
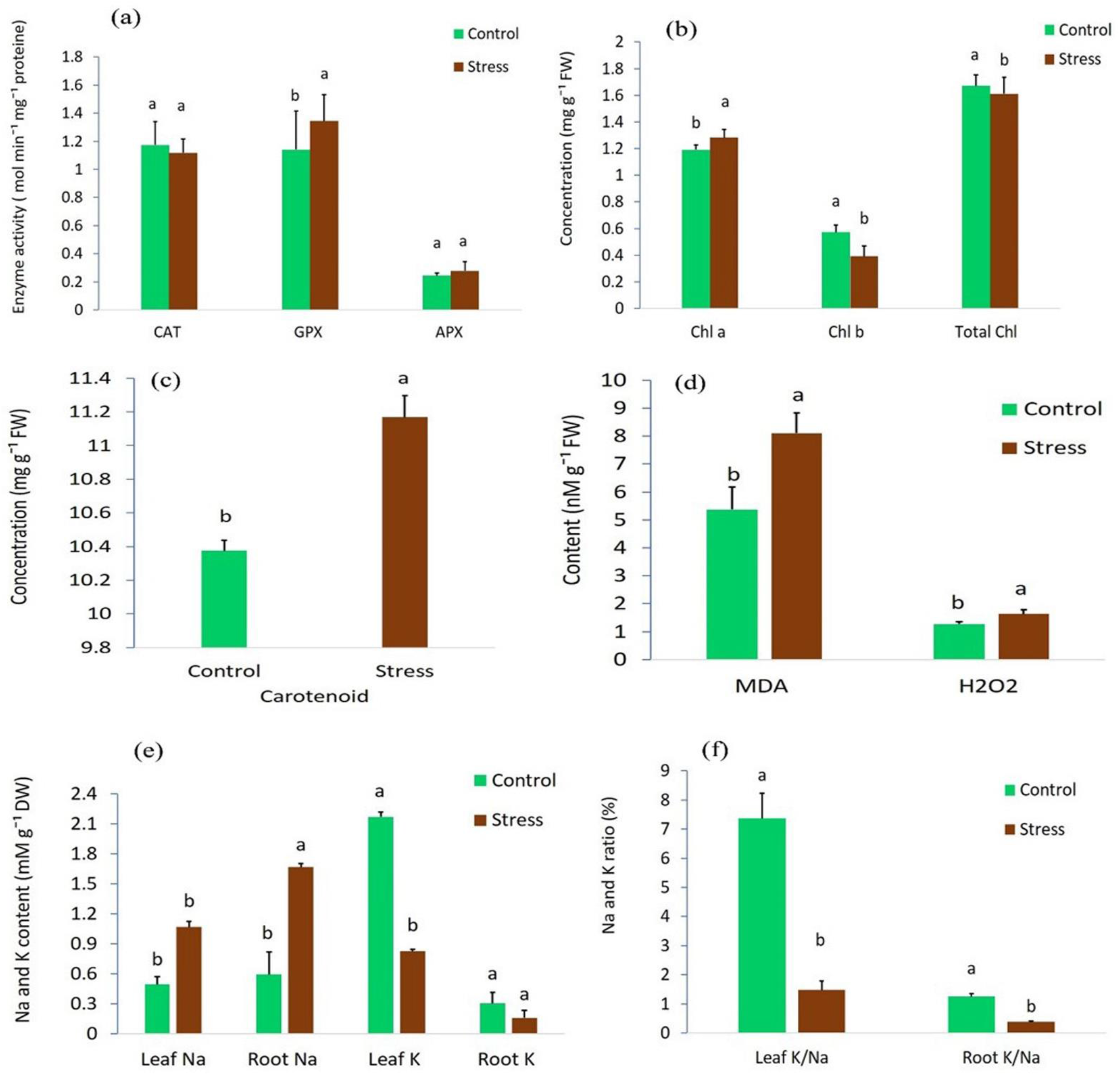
Effects of salt stress on (a) catalase (CAT), guaiacol peroxidases (GPX), ascorbate peroxidase (APX), (b) chlorophyll a (Chl_a_), chlorophyll b (Chl_b_) and total chlorophyll (Tchl) (c) carotenoid concentration, (d) MDA and H_2_O_2_ content, (e) root and leaf Na and K contents and (f) root and leaf Na/K ratios in parents (‘Roshan’, ‘Chinese spring’, *Ae. cylindrica*) and their F_1_ interspecific hybrids (amphidiploids). Bars represent means ± SE and bars with the same letter do not significantly differ at *p*<0.05.

Salt stress led not only to significant increases in leaf MDA from 5.37 to 8.10 nM g^−1^ FW but also to meaningful enhancements of 1.27 –1.63 mM g^−1^ FW in leaf H_2_O_2_ in the control and stressed plants, respectively (Fig. 3d). Moreover, its effect on MDA depended on genotype, with the highest observed in the female parents (namely, 115.9% in ‘Roshan’ and 149.30% in ‘Chinese Spring’ cultivars) (Table 4). An overall increase of 27.82% was observed in leaf H_2_O_2_ content of the genotypes tested (Fig. 3d), while much higher enhancements were observed in *Ae. cylindrica* and amphidiploid plants than in the female parents (Table 4).

Significant differences in leaf and root Na and K contents were observed among the genotypes under the control and salt-stress conditions (Table 1). Compared to the control treatment, salt stress was observed to raise Na contents in both leaf (from 0.49 to 1.07 mM g^−1^ DW) and root (from 0.59 to 1.51 mM g^−1^ DW) (Fig. 3e). In addition, salt stress caused a significant decrease in K content in both root (48%) and leaf (62%) tissues, with the leaf K content being evidently higher (Fig. 3e). Moreover, salt stress caused significantly decreased K/Na ratios in both root and leaf tissues (Fig. 3f). Nonetheless, the amphidiploid plants and *Ae. cylindrica* exhibited lower Na contents, higher K contents, and elevated K/Na ratios in both their root and leaf tissues than those measured for the wheat cultivars tested (Table 4). Compared to the female parents, the amphidiploids and wild parents exhibited higher Na accumulations in their roots than in their leaves (Table 4).

Fig. 4 shows the linear relationships between leaf dry matter (% control) and leaf mineral status (Na, K content, and K/Na ratio) under salt stress. Clearly, dry matter is significantly and negatively correlated with Na content (*r*^2^ =0.40, *p* < 0.01) (Fig. 4a) while it is significantly and positively correlated with K content (*r*^2^ = 0.58, *p* < 0.01) (Fig. 4b) and K/Na ratios (*r*^2^ =0.33, *p* < 0.01) (Fig. 4c). The genotypes superior with respect to dry matter such as the interspecific hybrid plants and *Ae. cylindrica* exhibited the lowest Na contents but higher K contents and K/Na ratios in their leaves when subjected to salt stress with 250 mM NaCl (Fig. 4).

**Fig 4:**
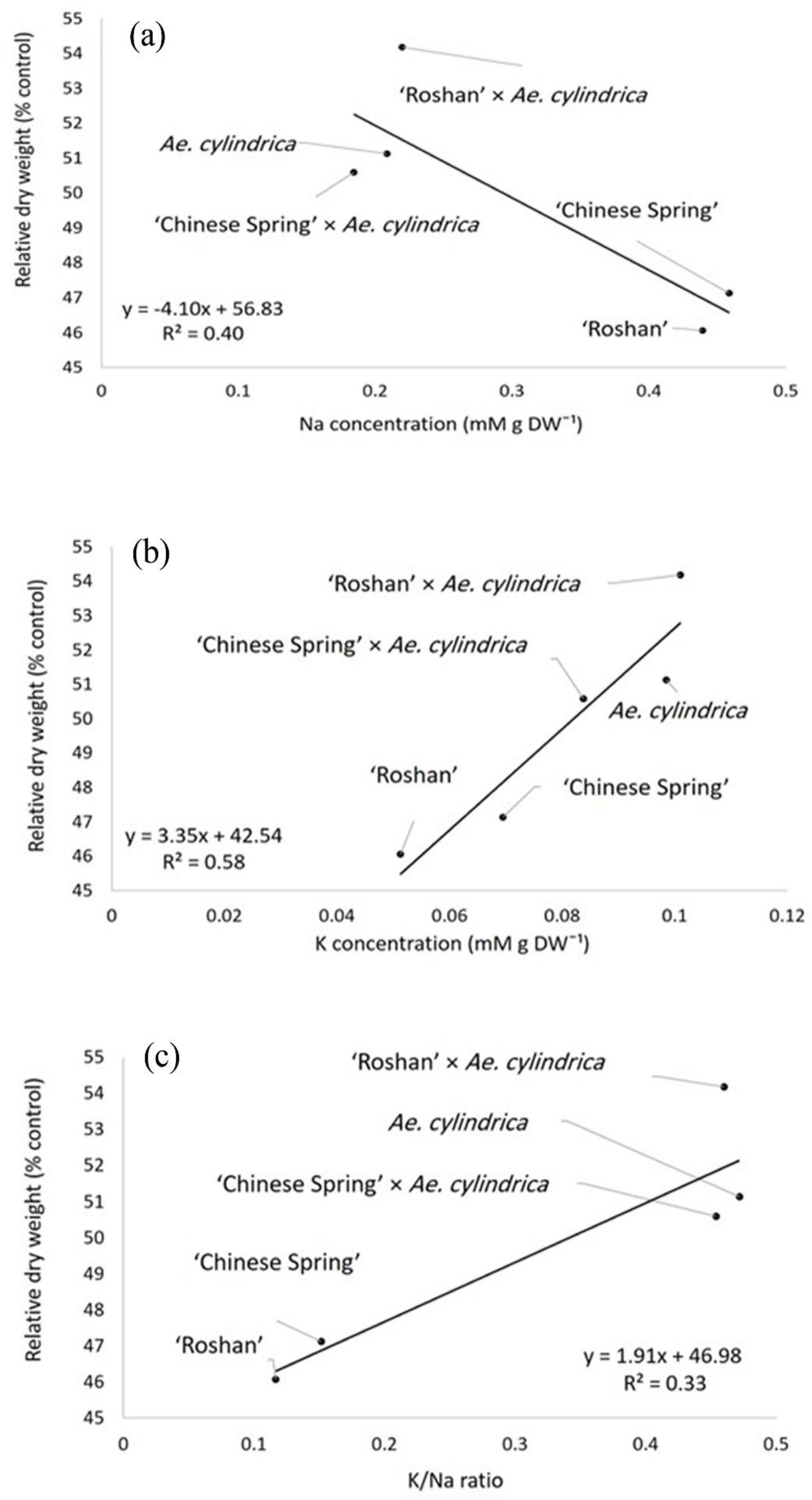
Relationship between dry matter (% control) and (a) leaf Na content, (b) leaf K content, and (c) leaf K/Na ratio of F_1_ interspecific hybrids and their parents in salt stress condition (250 mM NaCl).

Under the control conditions, FW and DW were positively correlated with Car, CAT, APX, and GPX activities as well as leaf K/Na ratio and root K content while they were negatively correlated with H_2_O_2_ content and leaf Na content. The correlation coefficients calculated for the data obtained under the 250 mM NaCl treatment showed that FW and DW were negatively correlated not only with MDA but with leaf and root Na contents as well but positively correlated with H_2_O_2_ content and leaf K/Na ratio as well as leaf and root K contents (Table 5). Meanwhile, a positive correlation was established between CAT activity and DW while H_2_O_2_ was at the same time significantly correlated with root K content as well as leaf and root K/Na ratios but negatively correlated with MDA and leaf Na content. Although leaf Na content showed negative correlations with APX and GPX activities under salt stress, leaf K content was positively associated with them. Finally, leaf and root K/Na ratios were found positively correlated (Table 5).

**Table 5:** Correlation coefficients of the traits studied under the control (above the diagonal) and salt-stress (250 mM NaCl) (below the diagonal) treatments

| Trait | FW | DW | Chl<sub>a</sub> | Chl<sub>b</sub> | Tchl | Car | CAT | APX | GPX |
| --- | --- | --- | --- | --- | --- | --- | --- | --- | --- |
| FW | 1 | 0.89<sup>*</sup> | 0.09<sup>ns</sup> | 0.34<sup>ns</sup> | 0.25<sup>ns</sup> | 0.62<sup>*</sup> | 0.70<sup>**</sup> | 0.52<sup>*</sup> | 0.55<sup>*</sup> |
| DW | 0.52<sup>*</sup> | 1 | 0.08<sup>ns</sup> | 0.40<sup>ns</sup> | 0.28<sup>ns</sup> | 0.68<sup>**</sup> | 0.56<sup>*</sup> | 0.53<sup>*</sup> | 0.51<sup>*</sup> |
| Chl<sub>a</sub> | 0.33<sup>ns</sup> | 0.19<sup>ns</sup> | 1 | 0.78<sup>**</sup> | 0.91<sup>**</sup> | 0.48<sup>ns</sup> | -0.16<sup>ns</sup> | 0.23<sup>ns</sup> | 0.13<sup>ns</sup> |
| Chl<sub>b</sub> | 0.15<sup>ns</sup> | 0.07<sup>ns</sup> | 0.61<sup>*</sup> | 1 | 0.96<sup>**</sup> | 0.74<sup>**</sup> | -0.04<sup>ns</sup> | 0.17<sup>ns</sup> | 0.24<sup>ns</sup> |
| Tchl | 0.26<sup>ns</sup> | 0.14<sup>ns</sup> | 0.88<sup>**</sup> | 0.92<sup>**</sup> | 1 | 0.67<sup>**</sup> | -0.07<sup>ns</sup> | 0.21<sup>ns</sup> | 0.21<sup>ns</sup> |
| Car | 0.12<sup>ns</sup> | 0.09<sup>ns</sup> | 0.93<sup>**</sup> | 0.72<sup>**</sup> | 0.91<sup>**</sup> | 1 | 0.17<sup>ns</sup> | 0.30<sup>ns</sup> | 0.45<sup>ns</sup> |
| CAT | 0.45<sup>ns</sup> | 0.51<sup>*</sup> | 0.27<sup>ns</sup> | 0.24<sup>ns</sup> | 0.29<sup>ns</sup> | 0.15<sup>ns</sup> | 1 | 0.26 | 0.44<sup>ns</sup> |
| APX | 0.52<sup>*</sup> | 0.17<sup>ns</sup> | -0.50<sup>ns</sup> | -0.39<sup>ns</sup> | -0.49<sup>ns</sup> | -0.44<sup>ns</sup> | -0.34<sup>ns</sup> | 1 | 0.34<sup>ns</sup> |
| GPX | 0.38<sup>ns</sup> | 0.25<sup>ns</sup> | -0.09<sup>ns</sup> | -0.36<sup>ns</sup> | -0.26<sup>ns</sup> | -0.27<sup>ns</sup> | -0.11<sup>ns</sup> | 0.20<sup>ns</sup> | 1 |
| H<sub>2</sub>O<sub>2</sub> | 0.54<sup>*</sup> | 0.52<sup>*</sup> | -0.41<sup>ns</sup> | -0.06<sup>ns</sup> | -0.24<sup>ns</sup> | -0.34<sup>ns</sup> | 0.05<sup>ns</sup> | 0.53<sup>*</sup> | 0.02<sup>ns</sup> |
| MDA | -0.74<sup>**</sup> | -0.52<sup>*</sup> | 0.16<sup>ns</sup> | -0.12<sup>ns</sup> | 0.06<sup>ns</sup> | -0.05<sup>ns</sup> | 0.32<sup>ns</sup> | -0.25<sup>ns</sup> | 0.47<sup>ns</sup> |
| Leaf Na | -0.83<sup>**</sup> | -0.52<sup>*</sup> | 0.19<sup>ns</sup> | 0.02<sup>ns</sup> | 0.11<sup>ns</sup> | -0.02<sup>ns</sup> | 0.18<sup>ns</sup> | -0.63<sup>*</sup> | -0.51<sup>*</sup> |
| Leaf K | 0.62<sup>*</sup> | 0.58<sup>*</sup> | -0.05<sup>ns</sup> | -0.01<sup>ns</sup> | -0.02<sup>ns</sup> | 0.11<sup>ns</sup> | -0.37<sup>ns</sup> | 0.52<sup>*</sup> | 0.59<sup>*</sup> |
| Leaf K/Na | 0.78<sup>**</sup> | 0.56<sup>*</sup> | -0.17<sup>ns</sup> | -0.10<sup>ns</sup> | -0.14<sup>ns</sup> | 0.05<sup>ns</sup> | -0.34<sup>ns</sup> | 0.59<sup>*</sup> | -0.47<sup>ns</sup> |
| Root Na | -0.57<sup>*</sup> | -0.64<sup>**</sup> | -0.02<sup>ns</sup> | 0.23<sup>ns</sup> | 0.13<sup>ns</sup> | 0.06<sup>ns</sup> | -0.34<sup>ns</sup> | -0.60<sup>*</sup> | -0.27<sup>ns</sup> |
| Root K | 0.81<sup>**</sup> | 0.53<sup>*</sup> | -0.18<sup>ns</sup> | -0.08<sup>ns</sup> | -0.13<sup>ns</sup> | -0.04<sup>ns</sup> | -0.33<sup>ns</sup> | 0.50<sup>ns</sup> | -0.16<sup>ns</sup> |
| Root K/Na | 0.38<sup>ns</sup> | 0.38<sup>ns</sup> | -0.02<sup>ns</sup> | -0.23<sup>ns</sup> | -0.15<sup>ns</sup> | -0.06<sup>ns</sup> | 0.09<sup>ns</sup> | 0.70<sup>**</sup> | 0.19<sup>ns</sup> |

| Trait | H<sub>2</sub>O<sub>2</sub> | MDA | Leaf Na | Leaf K | Leaf K/Na | Root Na | Root K | Root K/Na |
| --- | --- | --- | --- | --- | --- | --- | --- | --- |
| FW | -0.60<sup>*</sup> | -0.04<sup>ns</sup> | -0.89<sup>**</sup> | 0.26<sup>ns</sup> | 0.68<sup>**</sup> | 0.40<sup>ns</sup> | 0.63<sup>*</sup> | 0.24<sup>ns</sup> |
| DW | -0.68<sup>**</sup> | 0.11<sup>ns</sup> | -0.66<sup>**</sup> | 0.06<sup>ns</sup> | 0.61<sup>**</sup> | 0.29<sup>ns</sup> | 0.58<sup>*</sup> | 0.26<sup>ns</sup> |
| Chl<sub>a</sub> | -0.37<sup>ns</sup> | 0.10<sup>ns</sup> | 0.10<sup>ns</sup> | 0.13<sup>ns</sup> | -0.07<sup>ns</sup> | -0.39<sup>ns</sup> | 0.05<sup>ns</sup> | 0.25<sup>ns</sup> |
| Chl<sub>b</sub> | -0.52<sup>*</sup> | 0.41<sup>ns</sup> | 0.33<sup>ns</sup> | 0.23<sup>ns</sup> | -0.17<sup>ns</sup> | -0.20<sup>ns</sup> | 0.29<sup>ns</sup> | 0.35<sup>ns</sup> |
| Tchl | -0.49<sup>ns</sup> | 0.30<sup>ns</sup> | 0.25<sup>ns</sup> | 0.20<sup>ns</sup> | -0.11<sup>ns</sup> | -0.29<sup>ns</sup> | 0.18<sup>ns</sup> | 0.32<sup>ns</sup> |
| Car | -0.78<sup>**</sup> | 0.30<sup>ns</sup> | -0.49<sup>ns</sup> | 0.22<sup>ns</sup> | -0.27<sup>ns</sup> | -0.06<sup>ns</sup> | 0.47<sup>ns</sup> | 0.40<sup>ns</sup> |
| CAT | -0.27<sup>ns</sup> | 0.27<sup>ns</sup> | 0.67<sup>**</sup> | 0.27<sup>ns</sup> | -0.64<sup>**</sup> | 0.44<sup>ns</sup> | 0.18<sup>ns</sup> | -0.15<sup>ns</sup> |
| APX | -0.16<sup>ns</sup> | -0.31<sup>ns</sup> | 0.51<sup>*</sup> | 0.48<sup>ns</sup> | -0.12<sup>ns</sup> | 0.31<sup>ns</sup> | 0.47<sup>ns</sup> | 0.18<sup>ns</sup> |
| GPX | -0.46<sup>ns</sup> | -0.03<sup>ns</sup> | 0.52<sup>*</sup> | 0.44<sup>ns</sup> | 0.19<sup>ns</sup> | 0.17<sup>ns</sup> | 0.40<sup>ns</sup> | 0.21<sup>ns</sup> |
| H<sub>2</sub>O<sub>2</sub> | 1 | -0.12<sup>ns</sup> | -0.40<sup>ns</sup> | 0.03<sup>ns</sup> | 0.38<sup>ns</sup> | -0.18<sup>ns</sup> | -0.26<sup>ns</sup> | -0.06<sup>ns</sup> |
| MDA | -0.59<sup>*</sup> | 1 | -0.07<sup>ns</sup> | -0.46<sup>ns</sup> | -0.28<sup>ns</sup> | 0.02<sup>ns</sup> | -0.33<sup>ns</sup> | -0.29<sup>ns</sup> |
| Leaf Na | -0.60<sup>*</sup> | 0.73<sup>**</sup> | 1 | 0.49<sup>ns</sup> | -0.65<sup>**</sup> | 0.40<sup>ns</sup> | 0.57<sup>*</sup> | 0.22<sup>ns</sup> |
| Leaf K | 0.40<sup>ns</sup> | -0.79<sup>**</sup> | -0.75<sup>**</sup> | 1 | 0.32<sup>ns</sup> | 0.04<sup>ns</sup> | 0.66<sup>**</sup> | 0.60<sup>*</sup> |
| Leaf K/Na | 0.53<sup>*</sup> | -0.76<sup>**</sup> | -0.92<sup>**</sup> | 0.90<sup>**</sup> | 1 | -0.3<sup>ns</sup> | 0.05<sup>ns</sup> | 0.28<sup>ns</sup> |
| Root Na | -0.48<sup>ns</sup> | -0.07<sup>ns</sup> | 0.41<sup>ns</sup> | -0.08<sup>ns</sup> | -0.33<sup>ns</sup> | 1 | 0.42<sup>ns</sup> | -0.59<sup>*</sup> |
| Root K | 0.63<sup>*</sup> | -0.77<sup>**</sup> | -0.74<sup>**</sup> | 0.68<sup>**</sup> | 0.81<sup>**</sup> | -0.27<sup>ns</sup> | 1 | 0.71<sup>**</sup> |
| Root K/Na | 0.59<sup>*</sup> | -0.23<sup>ns</sup> | -0.60<sup>*</sup> | 0.35<sup>ns</sup> | 0.60<sup>*</sup> | -0.90<sup>**</sup> | 0.64<sup>*</sup> | 1 |

<sup>ns</sup> non-significant, <sup>\*</sup> $P < 0.05$ and <sup>\*\*</sup> $P < 0.01$

The results of PCA revealed that approximately 57% and 22% of the variance in data distribution could be explained by PC1 and PC2, respectively, under control condition (Fig. 5a) and 56% and 24% under salt-stress condition (Fig. 5b). Principal Component 1 (PC1) was found more closely correlated with FW DW, leaf K/Na ratio, and root K content as well as CAT and APX activities, but PC2 exhibited higher correlations with Chl_a_, Chl_b_, Tchl, and Car under the control conditions (Fig. 5a). Under salt stress, PC1 established negative correlations with H_2_O_2_, root K content, and K/Na ratio but it was positively associated with leaf Na content. On the other hand, PC2 was positively correlated with PDW but negatively correlated with root Na content. Ranking of the genotypes in the biplot identified amphidiploids and *Ae. cylindrica* as superior genotypes with respect to salt tolerance due to both their low PC1 and high PC2 values (Fig. 5b).

**Fig 5:**
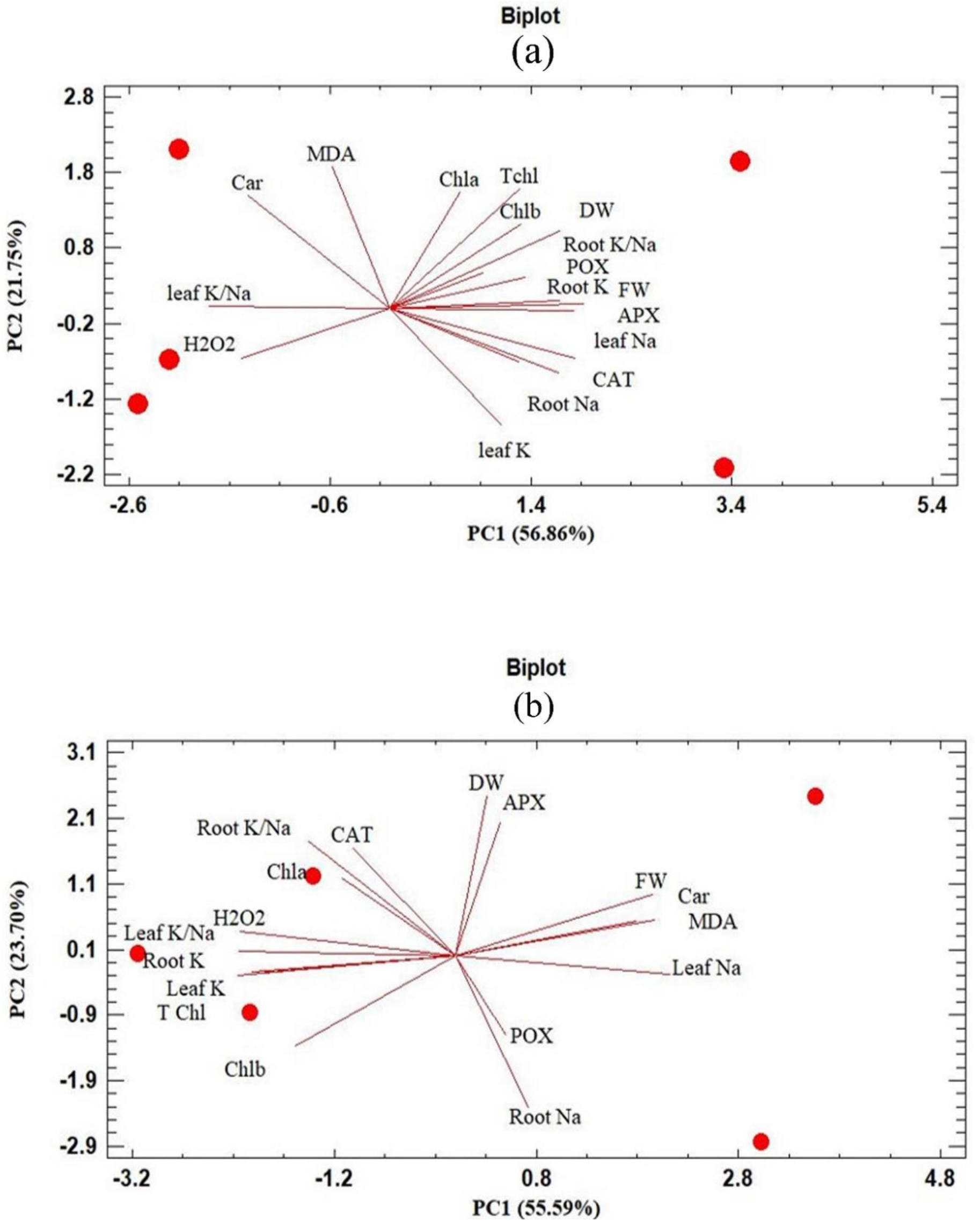
Bi-plot display of plant fresh weight (FW), plant dry weight (DW), chlorophyll a (Chl_a_), chlorophyll b (Chl_b_), total chlorophyll (Tchl), carotenoid (Car), catalase (CAT), ascorbate peroxidase (APX), guaiacol peroxidases (GPX), MDA and H_2_O_2_ content (nM g^−1^ FW), root and shoot Na and K contents DW^−1^, and K/Na ^+^ ratios (mM g) under (a) control and (b) 250 mM NaCl conditions.

### Expression of salt tolerance genes

The ANOVA results revealed the significant effects of salt stress on the expression profiles of *HKT1;5, NHX1* and *SOS1* in the roots and shoots (Table 6). More specifically, *HKT1;5* was induced by 2.6 times after 7 days of exposure to 250 mM NaCl as compared with the control (Fig. 6). Comparison of the expression patterns of the genes studied revealed that the expression level of the *NHX1* gene was by around six times higher than that under the control conditions (Fig. 7).

**Table 6:** Analysis of variance of the effects of NaCl treatments (0 and 250 mM at 7 days) on relative gene expression in plant tissues (shoot and root) in F_1_ interspecific hybrids and their parents

| Source of variation | df | Mean squares |  |  |
| --- | --- | --- | --- | --- |
|  |  | *HKT1;5* | *NHX1* | *SOS1* |
| Salt stress (S) | 1 | 14.79** | 58.70** | 5.07** |
| Generation (G) | 4 | 3.37** | 1.32** | 0.36** |
| Tissue (T) | 1 | 5.99** | 11.23** | 0.37** |
| S × G | 4 | 0.56** | 0.57** | 0.10** |
| S × T | 1 | 9.12** | 8.54** | 0.18* |
| G × T | 4 | 0.21** | 0.15<sup>ns</sup> | 0.01<sup>ns</sup> |
| S × G × T | 4 | 1.05** | 0.162<sup>ns</sup> | 0.01<sup>ns</sup> |
| Residual | 40 | 0.06** | 0.97 | 0.01 |
| CV (%) | - | 13.87* | 10.68 | 12.02 |

<sup>ns</sup> non-significant, \* $p < 0.05$ and \*\* $p < 0.01$

**Fig 6:**
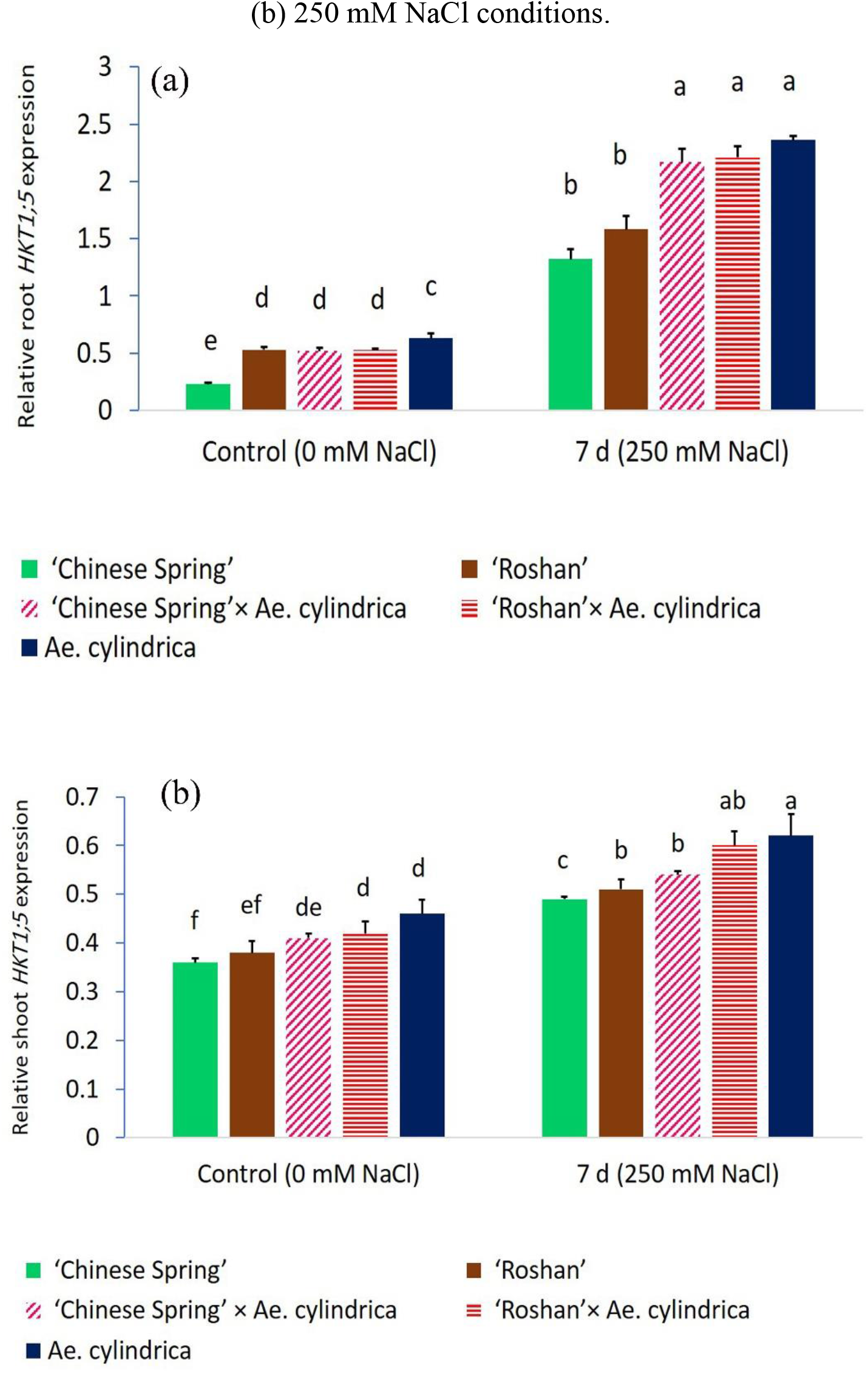
Expression analysis of the *HKT1;5* gene in root (a) and shoot (b) tissues of F_1_ interspecific hybrids and their parents. Bars represent means ± SE and bars headed by same letter are not significantly different at *p*<0.05.

**Fig 7:**
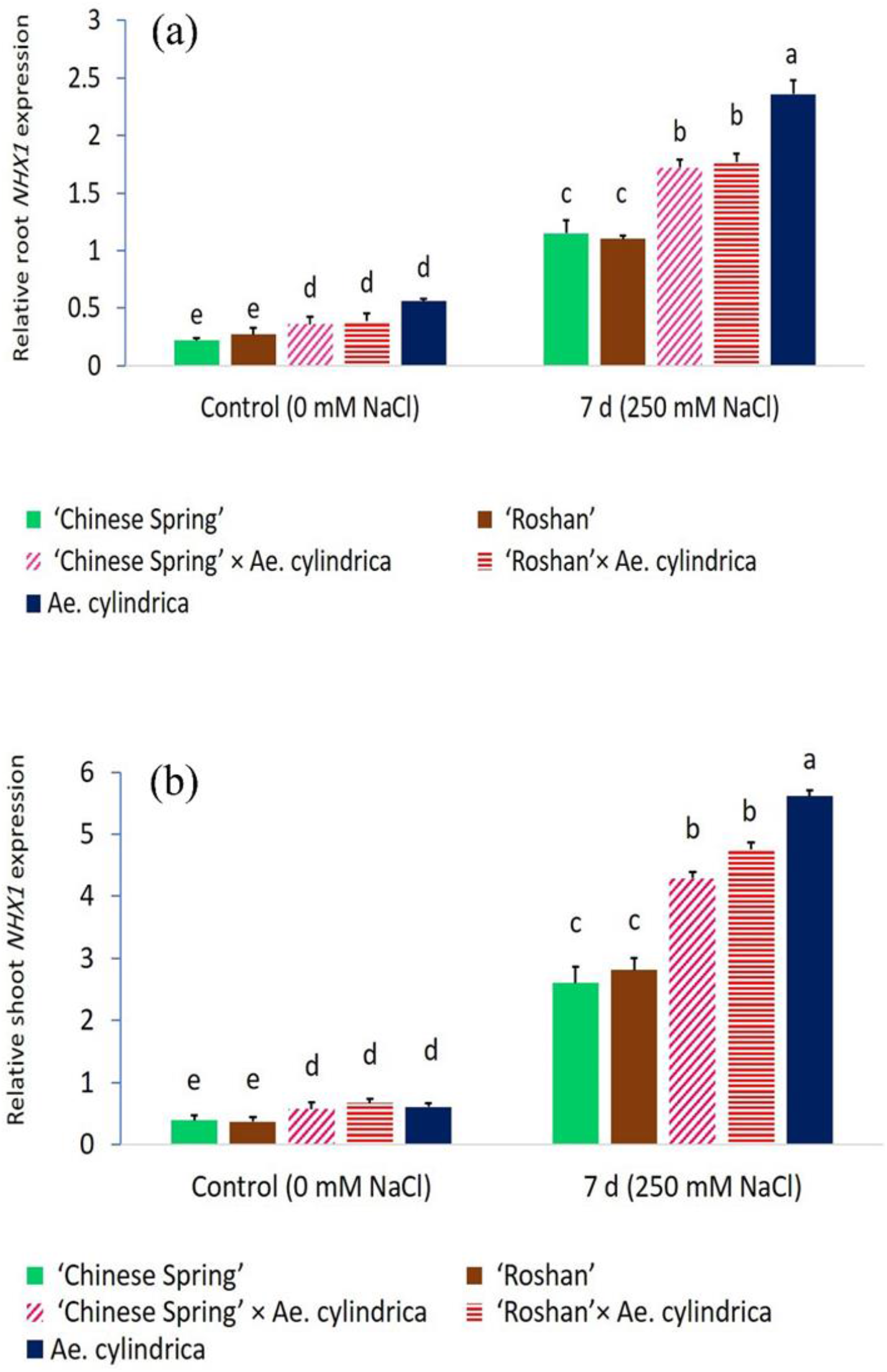
Expression analysis of the *NHX1* gene in root (a) and shoot (b) tissues of F_1_ interspecific hybrids and their parents. Bars represent means ± SE and bars headed by same letter are not significantly different at *p*<0.05.

At the tissue level, *HKT1;5* expressed mainly in the roots; indeed, its expression level in the root tissues was 2.4 times higher than that in the shoot tissues (Fig. 6). On the other hand, the transcript level of *NHX1* in the shoots was significantly greater (2.2-fold) than that in the roots (Fig. 7). Although, the transcripts of *SOS1* were upregulated in the root and shoot tissues of salt-stressed plants, the expression was slightly higher in the roots than the shoots (Fig. 8).

**Fig 8:**
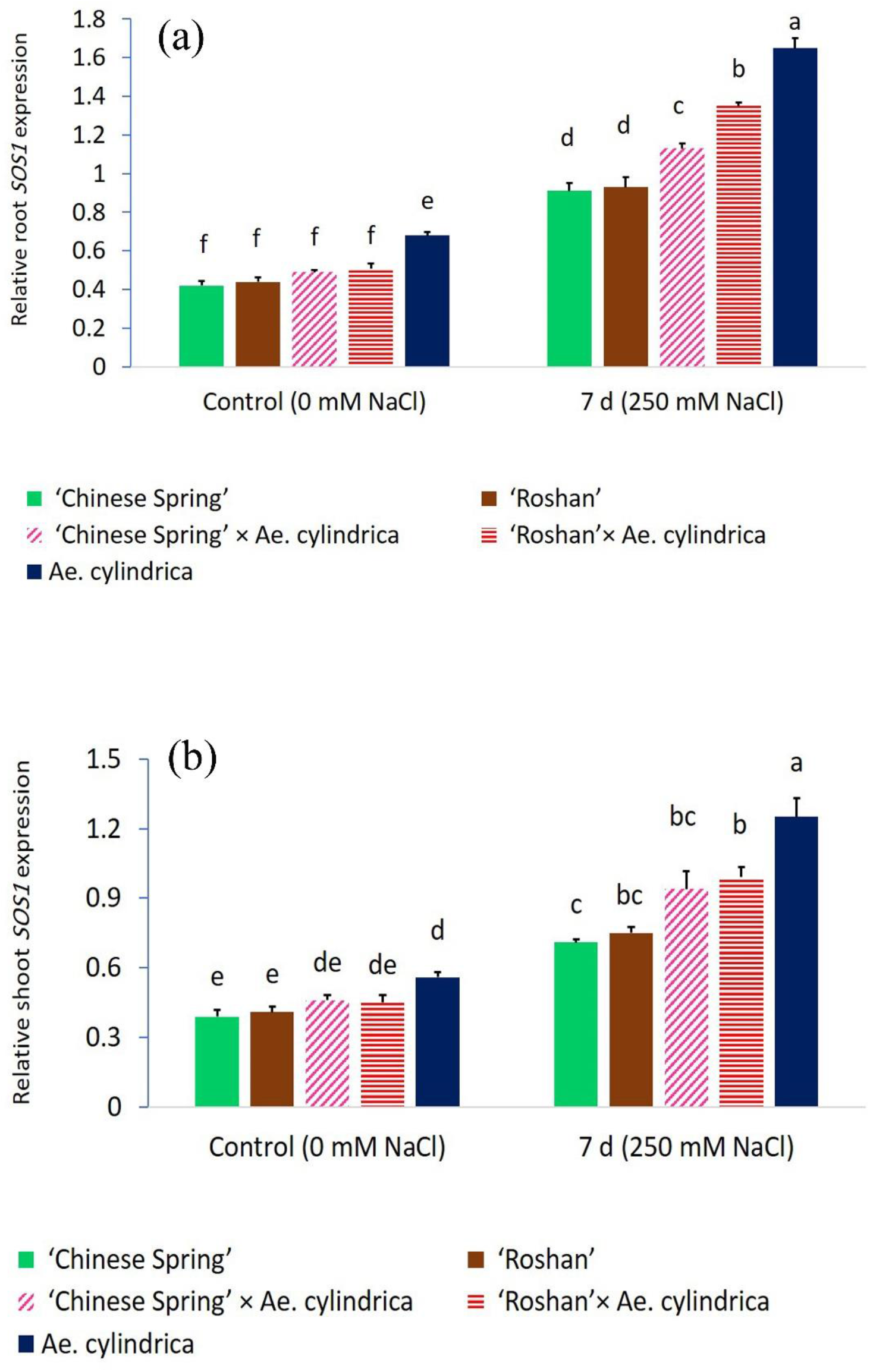
Expression analysis of the *SOS1* gene in root (a) and shoot (b) tissues of F_1_ interspecific hybrids and their parents. Bars represent means ± SE and bars headed by same letter are not significantly different at *p*<0.05.

The expression pattern analyses also showed significant differences in the responses of amphidiploids and their parents to the 250 mM NaCl stress (Table 6). Neither were any significant differences observed in the *HKT1;5* expression in the root tissues of *Ae. cylindrica* and amphidiploids plants, while these plants were significantly different in this respect from the wheat parents (Fig. 6a). *HKT1;5* transcript accumulations in the roots of *Ae. cylindrica* as well as ‘Roshan’ × *Ae. cylindrica*, and ‘Chinese Spring’ × *Ae. cylindrica* amphidiploids, respectively, under salt-stress were 3.7-fold, 4.2-fold, and 4.1-fold relative to those of the corresponding controls (Fig. 6a). Compared to ‘Chinese Spring’ and ‘Roshan’ cultivars, *Ae. cylindrica* and amphidiploids exhibited much higher *HKT1;5* expressions in the shoot. However, no significant differences were detected between the amphidiploids derived from ‘Roshan’ × *Ae. cylindrica* and ‘Chinese Spring’ × *Ae. cylindrica* crosses at the tissue level (Fig. 6b).

The *NHX1* transcripts of the salt-tolerant *Ae. cylindrica* in both tissues significantly exceeded those of the others. Interestingly, comparable mRNA expression levels were observed for *NHX1* in the roots and shoots of the amphidiploids of the ‘Roshan’ × *Ae. cylindrica* and ‘Chinese Spring’ × *Ae. cylindrica* crosses (Fig. 7a and b). Compared to the the control conditions, around 4-fold increases were detected in the root tissues of *Ae. cylindrica*, ‘Roshan’ × *Ae. cylindrica* and ‘Chinese Spring’ × *Ae. cylindrica* amphidiploids under NaCl stress (Fig. 7a). Finally, several-fold increases were detected in the *NHX1* transcript levels in the shoots of not only *Ae. cylindrica* but of the amphidiploids of ‘Roshan’ × *Ae. cylindrica* and ‘Chinese Spring’ × *Ae. cylindrica* crosses as well under stress condition when compared with the values under control condition (Fig. 7b).

The *SOS1* transcripts of the salt-tolerant *Ae. cylindrica* genotype in both tissues significantly exceeded those of other genotypes. Interestingly, comparable mRNA expression levels were observed for *SOS1* in the roots and shoots of the amphidiploids of the ‘Roshan’ × *Ae. cylindrica* and ‘Chinese Spring’ × *Ae. cylindrica* crosses (Fig. 8). *Ae. cylindrica* and amphidiploids exhibited a much higher expression of *SOS1* in the plant tissues (shoot and root), when compared to wheat cultivars (Fig. 8). However, Fig. 8 also shows that salt stress resulted in a significant increase in the *SOS1* transcript levels in the shoots and roots of all genotypes.

## Discussion

Salt stress is one of the most limiting factors in the production of such crop plants as wheat. Plant osmoregulation, protection against oxidative damage, and the strategy of maintaining a low level of cytosolic Na content – i.e., Na exclusion – intracellular compartmentation, and K/Na homeostasis are among the most important mechanisms of salt tolerance (Arzani 2008; Munns and Tester 2008). Wild relatives of wheat are valuable resources of tolerance to abiotic factors, whereby they have paved a long course of evolution via natural selection in the Fertile Crescent as their land of origin (Arzani and Ashraf 2016). *Aegilops* is the closest relative of the *Triticum* genus within the Triticeae tribe. The most salt-tolerant species from the 22 *Aegilops* species is *Ae. cylindrica*. Breeding would have relied primarily upon introgression of wild material, which offers a substantial improvement of salt tolerance in wheat.

In the present study, salt tolerance as well as the gene expression profiles of *HKT1;5* and *NHX1* were studied in the relevant amphidiploid plants derived from the ‘Chinese Spring’ × *Ae. cylindrica* and ‘Roshan’ × *Ae. cylindrica* crosses when subjected to salt stress with 250 mM NaCl. It was found that the amphidiploids derived from both interspecific crosses were less affected by salt stress than the wheat cultivars (i.e., the female parents) were and that they were either comparable or inferior to their male parent (*Ae. cylindrica*) with respect to the morphological and physiological traits or the gene expression profiles studied. Since availability of genetic resources of salt tolerance is an indispensable component of salt-tolerance improvement scheme (Arzani and Ashraf 2016), the synthesized amphiploids are potentially valuable resources for salt-tolerance improvement in common wheat through the backcrossing approach.

In the current study, both flag leaf area and plant height were found significantly influenced by genotype and salt stress. These findings are in line with the assumption that genotypes with smaller flag leaf areas and plant heights possess lower yield losses under salt-stress conditions (Chamekh et al. 2016) as a result of low accumulations of toxic ions in their photosynthetically active leaves (Pan et al. 2020). Although salt stress was observed to cause a marked reduction in the growth of both wheat cultivars (‘Chinese Spring’ and ‘Roshan’), the interspecific hybrid plants exhibited significantly lower reductions as did *Ae. cylindrica* as well. Salt stress is claimed not only to have an important impact on plant biomass but also to stand as a key trait for assessing salt tolerance in crop plants (Munns et al. 2016; Negrao et al. 2017). Studying barley, Han et al. (2018) found tolerant genotypes to exhibit lower reductions in their dry biomass and less Na^+^ accumulated in their roots and shoots than did the other wild or cultivated genotypes investigated. Consistent with our data, Zeeshan et al. (2020) recently reported that the reduction in plant dry weight was much more noticeable in sensitive wheat cultivars subjected to salt stress. Similar results had also been reported by Kiani et al. (2015) and Kumar et al. (2017).

Although plants are potentially capable of escaping salt, there might exist an evolutionary trade-off between salt escape and other salt tolerance strategies. Our findings confirm that while the phonological traits (DH, DP, and DM) declined under salt stress, the amphidiploid plants and *Ae. cylindrica*, with already low growth periods under both the control and salt-stress conditions, exhibited less plasticity. Indeed, shortened life cycle is one of the important strategies that wild species more often use to cope with drought, salt, and temperature stresses. Phenological stages in the current study were observed to occur earlier in response to salt stress, which is consistent with the observations reported by Chamekh et al. (2016). Phenotypically plastic genotypes that adjust phenology through differential gene expressions following salt-stress cues are maintained to offer the valuable germplasm for the salt escape mechanism (Munns et al. 2016).

The superior salt tolerance of the amphidiploids and *Ae. cylindrica* was substantiated by a non-significant decline in their Tchl in contrast to the significant Tchl reduction observed in wheat cultivars. The deleterious effects of salt stress on chlorophyll content in wheat have also been reported elsewhere (Rauf et al. 2010; Kumar et al. 2017; Al-Ashkar et al. 2019).

Significant increases in CAT, APX, and GPX activities were detected in the amphidiploids of both the ‘Roshan’ × *Ae. cylindrica* and Chinese Spring ×*Ae. cylindrica* crosses while they were more or less declining in their female parents due to the stress (Table 2). Increasing antioxidant enzyme activities such as those of APX, GPX, and CAT provide further support for the above inferences about superior salt tolerance of the amphidiploids over that of the wheat cultivars. In addition, CAT was observed to have a positive correlation with DW, a finding that is in agreement with recent empirical results reported by Al-Ashkar et al. (2019). Scavenging reactive oxygen species or free radicals (namely, O_2^−^_, OH^−^, and H_2_O_2_) that are extremely damaging oxidizing agents are modulated by superoxide dismutase (SOD) to remove the toxicity of the superoxide anion. The ROS produced in chloroplasts are reduced to H_2_O_2_and O_2_ by SOD; the H_2_O_2_ and its derivatives thus produced are rapidly removed or reduced by APX or GPX (Sharma et al. 2016).

The genotypes investigated in this study showed differential responses to the discriminate K and Na in the soil solution. In addition, a higher Na accumulation was detected in the roots than in the leaves under salt stress conditions. The discriminative dynamic features to uptake K^+^ and Na^+^ from the soil solution, the preferential accumulation of K^+^, and the exclusion of Na^+^ are known as the important mechanisms involved in the salt tolerance of the tribe Triticeae (Shabala et al. 2013; Kumar et al. 2015; Zhu et al. 2016). Generally speaking, it is the goal of wheat breeders in saline-prone regions to develop salt-tolerant cultivars highly capable of Na^+^ exclusion, a trait that is found in such wild halophytically close progenitors of wheat as *Aegilops cylindrica* (Colmer et al. 2006; Arabbeigi et al. 2014).

Although salt stress elicited a decline in both root and leaf K contents, K accumulation in the leaves was by several folds higher in the roots under both salt stress and control conditions. However, the amphidiploids and *Ae. cylindrica* contained lower Na but higher K contents and K/Na ratios in both roots and leaves. These findings led the present authors to hypothesize that the amphidiploid plants exploited such superior features as diminishing Na influx into the root, Na retrieval from the xylem, regulating Na^+^ efflux from the root, Na xylem loading, Na recirculation in the phloem, intracellular compartmentation of Na into the vacuoles, and Na secretion from the leaf (Arzani and Ashraf 2016; Kronzucker and Britto 2011; Shabala 2011; Shabala et al. 2016). In addition, root Na and K content were found in the current study to be correlated positively and negatively, respectively, with DW under salt stress conditions, suggesting the critical role of the root in Na^+^ transport and ion homeostasis. Plant root perpetually strives to adapt itself to perfect conformity with the saline environment through multiple strategies, one of which is conceding stress-elicited K^+^ efflux for signaling commitments without endangering the cell via programmed cell death (PCD) (Rubio et al. 2020; Shabala et al. 2015). Moreover, PCD is also reportedly induced under salt stress conditions by SKOR and GORK-mediated K^+^ efflux (Arzani 2018).

The highest increase in MDA/lipid peroxidation was observed in the wheat cultivars (namely, ‘Chinese Spring’ and ‘Roshan’) while the lowest increase was recorded for the amphidiploids and *Ae. cylindrica* (Table 2). Consistent with our findings, lower levels of lipid peroxidation had been reported in salt-tolerant *Ae. cylindrica* genotypes (Kiani et al. 2015) and salt-tolerant wheat cultivars (Kumar et al. 2017). These findings are also complements to those reported on wheat and barley by Zeeshan et al. (2020) who found that salt-stress induced significant changes in MDA content among their cultivars, with the highest recorded for a sensitive (Sunmate) wheat cultivar. MDA is one of the end products of lipid peroxidation indicating the status of ROS (reactive oxygen species). ROS appear to act as signaling molecules on the one hand, while excessive generation of ROS leads to oxidative damages to such fundamental biological molecules as proteins, RNA, DNA, and membranes (lipid peroxidation) as well as PCD, on the other (Arzani 2018).

The results of the current study showed that, similar to lipid peroxidation, H_2_O_2_ increased as a result of NaCl stress; this was accompanied by lower H_2_O_2_ accumulation in the amphidiploids and the hyper salt-tolerant genotype *Ae. cylindrica*. These results are underpinned by previous reports on the effects of salt stress on H_2_O_2_ accumulation in *Ae. cylindrica* and wheat genotypes (Sairam et al. 2005; Kiani et al. 2015). H_2_O_2_ plays an important role in the process of signaling salt-stress induced K^+^ efflux from plant cells through non-selective cation (NSCC) channels (Rubio et al. 2020). Furthermore, H_2_O_2_ reportedly modulates the expressed SKOR, identifying this K^+^ channel as a potential target for ROS signaling (Wu et al. 2018). Finally, enhanced levels of H_2_O_2_ trigger PCD to result in DNA laddering, caspase-3-like protein recruitment, and increased PARP cleavage (Arzani 2018).

Principal component analysis (PCA) serves to simplify a complex dataset of multiple dimensions by coalescing a set of variables into a few principal components (PCs) that capture the highest variance in the data. The relationships among the attributes are then used as efficient screening criteria. According to the biplot analysis of PC1 and PC2 in the current study, the amphidiploids and *Ae. cylindrica* (the male parent) were identified as superior salt tolerant genotypes. Examination of whole data also strengthened the claim that the amphidiploids are superior in terms of their salt tolerance to the wheat cultivars used as female parents.

HKTs and NHXs synergistically regulate Na^+^ exclusion in salt-tolerant genotypes. HKTs regulate Na^+^ homeostasis by controlling Na^+^ uptake into the root (Munns et al. 2006) and by excluding excessive Na^+^ from photosynthetic tissues (Arzani and Ashraf 2016). The hyper salt-tolerant USL26 genotype of *Ae. cylindrica* has been similarly reported to possess a high capability for Na^+^ exclusion conferred by C and D genomes (Arabbeigi et al. 2014). In the present study, this hyper salt-tolerant genotype was used to develop the amphidiploid plants. The expression analysis of *HKT1;5* and *NHX1* in the tested genotypes revealed that they could serve as potential candidates for Na^+^ exclusion from the roots and shoots of ‘Roshan’ × *Ae. cylindrica* and ‘Chinese Spring’ × *Ae. cylindrica* amphidiploids and their male parent. The *HKT1;5* and *NHX1* genes exhibited differential expression patterns in the shoot and the root such that *HKT1;5* had a higher expression in the roots than in the leaves but the opposite was true for *NHX1*.

Alteration in plant gene expression is induced by environmental stresses such as salt stress through either epigenetic changes or modulation of the core signaling networks that successively trigger a downstream network of interacting hormones and transcriptional regulators. The whole process leads to the adjustment of plant growth and development, generating molecular, physiological, and phenotypic plasticity in the plant. Previous study has demonstrated that *AtHKT1* facilitates Na^+^ homeostasis and modulates K^+^ nutrient status in the plant (Rus et al. 2004; Oyiga et al. 2018; Zhang et al. 2017). In *T. aestivum, HKT1;5* as the Na^+^ selective transporter plays an important role in improving salt tolerance by Na^+^ exclusion from the xylem vessels to its parenchyma cells (XPCs), thereby maintaining a higher K^+^/Na^+^ ratio and reducing leaf Na^+^ content (Oyiga et al. 2018; Rus et al. 2004). The results of the present study indicated that *HKT1;5* gained a higher expression in both the roots and shoots under the 250 mM NaCl treatment when compared with that under the control. Interestingly, the gene was to a significant level more strongly expressed in the roots than in the shoots. These findings are consistent with those previously reported, confirming the claimed significance of the genotype-and tissue-dependent patterns in the expression of *HKT1;5* in both wheat (Zamani et al. 2013) and *Ae. cylindrica* (Arabbeigi et al. 2018). The tissue-specific patterns observed can most likely be explained with recourse to the fact that *HKT1;5-A* encodes a plasma membrane Na^+^ transporter expressed in the root cells surrounding xylem vessels located at an ideal site for extracting Na^+^ from the xylem to reduce Na^+^ transfer to the leaves (Munns et al. 2012). *HvHKT1* exhibits a similar function in barley, in which the mRNA of the *HvHKT1* gene is chiefly transcribed in response to salt stress in the roots rather than in the shoots as evidenced by its expression at the plasma membrane of the root cells (stele and epidermis), which is the layer surrounding the xylem (Han et al. 2018).

It is a reasonable premise that the *SOS1* gene plays an important role in Na^+^ homeostasis, by removing Na^+^ out of the cytosol into the apoplastic space (Shi et al. 2002). The results of the current study support the findings of previous study in *Ae. cylindrica* (Arabbeigi et al. 2018), and *Puccinellia tenuiflora* (Zhang et al. 2017), and confirm the independent and joint contribution of the three genes (*NHX1, HKT1;5*, and *SOS1*) to salt stress tolerance.

In contrast to the roots, the shoots exhibited a much higher expression of *NHX1* where it increased more consistently under the 250 mM NaCl treatment. These findings, together with the higher Na accumulation observed in the roots of the amphidiploids and *Ae. cylindrica* genotype as well as the lower root Na content of the wheat cultivars under the NaCl treatment, suggest that certain mechanisms may be involved in promoting Na exclusion from the photosynthetic tissue in *Ae. cylindrica* and its interspecific hybrid progeny. Our results agree well with those of Nawaz et al. (2014) who reported that *NHX1* was more strongly expressed in the shoots of halophytic and glycophytic Brassicaceae species than in their roots. Experimenting with *Suaeda salsa*, however, Ma et al. (2004) reported that the expression of *SsNHX1* was enhanced by NaCl treatment across the entire plant tissues including its shoots and roots.

The results of the current study indicated that *NHX1* recorded a higher expression in the genotypes studied than did *HKT1;5*. Although salt exclusion from plant tissues is indisputably of utmost importance in handling salt injury, the salt tolerance of tissues is also a critical element of the plant’s overall salt tolerance.

## Conclusion

The different aspects of the current study lead to the general finding that amphidiploids exhibit more resemblance to their wild parent (a hyper-salt tolerant genotype of *Ae. cylindrica*). More specifically, we might draw the following conclusions from the data obtained. 1) Compared to the female parents, the interspecific hybrid plants and their wild parent were found less affected by salt stress in terms of growth, physiological, and phenological traits; 2) generally speaking, an interplay seems to exist between *HKT1;5, NHX1*, and *SOS1* which restrict root-to-shoot transport of Na and elevate the vacuolar sequestration of Na in the shoot tissues, respectively, thereby synergistically regulating Na^+^ homeostasis in *A. cylindrica* and amphidiploid plants subjected to salt stress; 3) the higher root and leaf K levels and K/Na ratios are associated with the plant’s higher salt tolerance; and 4) the synthesized amphiploids might potentially serve as valuable resources for improving salt tolerance in bread wheat by further backcrossing with *T. aestivum* L.

## Acknowledgements

This research was supported by funds from the Iran National Science Foundation (INSF) under Grant No. 96009225. Part of this research was conducted while the first author was on a study leave at the National Institute of Genetic Engineering and Biotechnology, Tehran, Iran. The final editing of the English manuscript was performed by Dr. Ezzatollah Roustazadeh from ELC, IUT.

## Author contribution statement

AA conceived and designed the research. The experiments were conducted by RK under the supervision of AA. SAMMM, and M.R. except the molecular analysis that was carried out under the supervision of KR. Finally, data analysis was performed by RK who also wrote down the draft of the manuscript with subsequent contributions from all the authors.

## References

Aebi H (1984) Catalase in vitro. Methods Enzymol 105: 121–126

Al-Ashkar I, Alderfasi A, El-Hendawy S, Al-Suhaibani N, El-Kafafi S, Seleiman MF (2019) Detecting salt tolerance in doubled haploid wheat lines. Agronomy 9: 211

Apse MP, Sottosanto JB, Blumwald E (2003) Vacuolar cation/H^+^ exchange, ion homeostasis, and leaf development are altered in a T-DNA insertional mutant of AtNHX1, the Arabidopsis vacuolar Na^+^/H^+^ antiporter. Plant J 36: 229–239

Arabbeigi M, Arzani A, Majidi MM, Kiani R, SayedbTabatabaei BE, Habibi F (2014) Salinity tolerance of Aegilops cylindrica genotypes collected from hyper-saline shores of Uremia Salt Lake using physiological traits and SSR markers. Acta Physiol Plant 36: 2243–2251

Arabbeigi M, Arzani A, Majidi MM, Sayed-Tabatabaei BE, Saha P (2018) Expression pattern of salt tolerance-related genes in Aegilops cylindrica. Physiol Mol Biol Plant 24: 61–73

Arino-Estrada G, Mitchell GS, Saha P, Arzani A, Cherry SR, Blumwald E, Kyme AZ (2019) imaging salt uptake dynamics in plants using PET. Sci Rep 9: 1–9

Arzani A (2008) Improving salinity tolerance in crop plants: a biotechnological view. In Vitro Cell Dev Biol Plant 44: 373–383

Arzani A (2018) Engineering programmed cell death pathways for enhancing salinity tolerance in crops. In: Kumar V., Wani S.H., Suprasanna P. and Tran LPS. (eds) Salinity Responses and Tolerance in Plants, Volume 2. Springer, pp 93–118

Arzani A, Ashraf M (2016) Smart engineering of genetic resources for enhanced salinity tolerance in crop plants. Crit Rev Plant Sci 35: 146–189

Arzani A, Darvey NL (2001) The effect of colchicine on triticale anther-derived plants: Microspore pre-treatment and haploid-plant treatment using a hydroponic recovery system. Euphytica 122 235–241

Bradford MM (1976) A rapid and sensitive method for the quantitation of microgram quantities of protein utilizing the principle of protein-dye binding. Anal Biochem 72: 248–254

Chamekh Z, Ayadi S, Karmous C, Trifa Y, Amara H, Boudabbous K, Yousfi S, Serret MD, Araus JL (2016) Comparative effect of salinity on growth, grain yield, water use efficiency, δ^13^C and δ^15^N of landraces and improved durum wheat varieties. Plant Science 251: 44–53

Colmer TD, Flowers TJ, Munns R (2006) Use of wild relatives to improve salt tolerance in wheat. J Exp Bot.57: 1059–1078

Demidchik V, Shabala SN, Davies JM (2007) Spatial variation in H_2O2_response of Arabidopsis thaliana root epidermal Ca^2+^ flux and plasma membrane Ca^2+^ channels. Plant J 49: 377–386

Duan HR, Ma Q, Zhang JL, Hu J, Bao AK, Wei L, et al (2015) The inward-rectifying KC channel SsAKT1 is a candidate involved in KC uptake in the halophyte Suaeda salsa under saline condition. Plant Soil 395: 173–187

Flowers TJ, Colmer TD (2015) Plant salt tolerance: adaptations in halophytes. Ann Bot 115: 327–331

Han Y, Yin S, Huang L, Wu X, Zeng J, Liu X, Zhang G (2018) A sodium transporter HvHKT1; 1 confers salt tolerance in barley via regulating tissue and cell ion homeostasis. Plant Cell Physiol 59: 1976–1989

Herzog V, Fahimi H (1973) Determination of the activity of peroxidase. Anal Biochem 55: 554–562

Houshmand S, Arzani A, Maibody SAM, Feizi M (2005) Evaluation of salt-tolerant genotypes of durum wheat derived from in vitro and field experiments. Field Crops Res 91: 345–354

Kiani R, Arzani A, Habibi F (2015) Physiology of salinity tolerance in Aegilops cylindrica. Acta Physiol Plant 37: 135

Kronzucker HJ, Britto DT (2011) Sodium transport in plants: a critical review. New Phytol 189: 54–81

Kumar S, Beena AS, Awana M, Singh A (2017) Physiological, biochemical, epigenetic and molecular analyses of wheat (Triticum aestivum) genotypes with contrasting salt tolerance. Front Plant Sci 8: 1151

Kumar M, Hasan M, Arora A, Gaikwad K, Kumar S, Rai RD, Singh A (2015) Sodium chloride-induced spatial and temporal manifestation in membrane stability index and protein profiles of contrasting wheat (Triticum aestivum L.) genotypes under salt stress. Ind J Plant Physiol 20: 271–275

Lichtenthaler HK, Buschmann C (2001) Chlorophylls and Carotenoides: measurement and characterization by UVVIS spectroscopy. John Wiley and Sons, Inc, New York

Livak KJ, Schmittgen TD (2001) Analysis of relative gene expression data using real-time quantitative PCR and the 2-DDCT method. Methods 25: 402–408

Ma XL, Zhang Q, Shi HZ, Zhu JK, Zhao YX, Ma CL, Zhang H (2004) Molecular cloning and different expression of a vacuolar Na^+^/H^+^ antiporter gene in Suaeda salsa under salt stress. Biol Plant 48: 219–225

Maathuis FJ, Ahmad I, Patishtan J (2014) Regulation of Na^+^ fluxes in plants. Front Plant Sci 5: 467

Munns R, James RA, Lauchli A (2006) Approaches to increasing the salt tolerance of wheat and other cereals. J Exp Bot 57: 1025–1043

Munns R, James RA, Xu, B, Athman A, Conn SJ, Jordans C, Plett D, Byrt CS, Hare RA, Tyerman SD, Tester M, Plett D, Gilliham M (2012) Wheat grain yield on saline soils is improved by an ancestral Na^+^ transporter gene. Nat Biotechnol 30: 360–364

Munns R, James RA, Gilliham M, Flowers TJ, Colmer TD (2016) Tissue tolerance: an essential but elusive trait for salt-tolerant crops. Funct Plant Biol 43: 1103–1113

Munns R, Tester M (2008) Mechanisms of salinity tolerance. Ann. Rev. Plant Biol 59: 651–681

Murray MG and Thompson WF (1980) Rapid isolation of high-molecular-weight plant DNA. Nucleic Acids Res 8: 4321–4325.

Nakano Y, Asada K (1981) Hydrogen peroxide is scavenged by ascorbate-specific peroxidase in spinach chloroplasts. Plant Cell Physiol 22: 867–880

Nawaz I, Iqbal M, Hakvoort HW, Bliek M, de Boer B, Schat H (2014) Expression levels and promoter activities of candidate salt tolerance genes in halophytic and glycophytic Brassicaceae. Environ Exp Bot 99: 59–66

Negrao S, Schmockel SM, Tester M (2017) Evaluating physiological responses of plants to salinity stress. Ann Bot 119: 1–11

Nemeth C, Yang CY, Kasprzak P, Hubbart S, Scholefield D, Mehra S, King J (2015) Generation of amphidiploids from hybrids of wheat and related species from the genera Aegilops, Secale, Thinopyrum, and Triticum as a source of genetic variation for wheat improvement. Genome 58: 71–79

Pan T, Liu M, Kreslavski VD, Zharmukhamedov SK, Nie C, Yu M, Kuznetsov VV, Allakhverdiev SI, Shabala S (2020) Non-stomatal limitation of photosynthesis by soil salinity. Crit Rev Environ Sci Technol 1–35, DOI: 10.1080/10643389.2020.1735231

Rauf S, Adil MS, Naveed A, Munir H (2010) Response of wheat species to the contrasting saline regimes. Pak J Bot 42: 3039–3045

Rubio F, Nieves-Cordones M, Horie T, Shabala S (2020) Doing ‘business as usual’ comes with a cost: evaluating energy cost of maintaining plant intracellular K^+^ homeostasis under saline conditions. New Phytol 225: 1097–1104

Rus A, Lee BH, Munoz-Mayor, A, Sharkhuu A, Miura K, Zhu JK, Hasegawa PM (2004) AtHKT1 facilitates Na^+^ homeostasis and K^+^ nutrition in planta. Plant Physiol 136: 2500– 2511

Oyiga BC, Sharma RC, Baum M, Ogbonnaya FC, Leon J, Ballvora A (2018) Allelic variations and differential expressions detected at quantitative trait loci for salt stress tolerance in wheat. Plant Cell Environ 41: 919–935

Sairam RK, Srivastava GC, Agarwal S, Meena RC (2005) Differences in antioxidant activity in response to salinity stress in tolerant and susceptible wheat genotypes. Biol Plant 49: 85–91

SAS Institute (2011) Base SAS 9.3 procedures guide. SAS Institute Inc., Cary, NC

Shabala L, Zhang J, Pottosin I et al (2016) Cell-type specific H^+^-ATPase activity enables root K^+^ retention and mediates acclimation to salinity. Plant Physiol 172: 2445–2458

Shabala S (2017) Signaling by potassium: another second messenger to add to the list? J Exp Bot 68: 4003–4007

Shabala S (2011) Physiological and cellular aspects of phytotoxicity tolerance in plants: the role of membrane transporters and implications for crop breeding for waterlogging tolerance. New Phytol 190: 289–298

Shabala S, Hariadi Y, Jacobsen SE (2013) Genotypic difference in salinity tolerance in quinoa is determined by differential control of xylem Na^+^ loading and stomatal density. J Plant Physiol 170: 906–914

Shabala S, Pottosin I (2014) Regulation of potassium transport in plants under hostile conditions: implications for abiotic and biotic stress tolerance. Physiol Plant 151: 257–279

Shabala S, Wu H, Bose J (2015) Salt stress sensing and early signaling events in plant roots: current knowledge and hypothesis. Plant Sci 241:109–119

Sharma NYS (2016) Reactive oxygen species, oxidative stress and ROS scavenging system in plants. J Chem Pharm Res 8 595–604

Shi H, Quintero FJ, Pardo JM, Zhu JK (2002) The putative plasma membrane Na^+^/H^+^ antiporter SOS1 controls long-distance Na^+^ transport in plants. Plant Cell 14: 465–477.

Taulavuori E, Hellstrom EK, Taulavuori K, Laine K (2001) Comparison of two methods used to analyse lipid peroxidation from Vaccinium myrtillus (L.) during snow removal, reacclimation and cold acclimation. J Exp Bot 52: 2375–2380

Velikova V, Yordanov I, Edreva A (2000) Oxidative stress and some antioxidant systems in acid rain treated bean plants. Protective role of exogenous polyamines. Plant Sci 151: 59– 66

Wu H, Zhang X, Giraldo JP, Shabala S (2018) It is not all about sodium: revealing tissue specificity and signalling roles of potassium in plant responses to salt stress. Plant Soil 431: 1–17

Yan K, Shao H, Shao C, Chen P, Zhao S, Brestic M, et al (2013) Physiological adaptive mechanisms of plants grown in saline soil and implications for sustainable saline agriculture in coastal zone. Acta Physiol Plant 35: 2867–2878

Zamani BM, Niazi A, Moghadam AA, Deihimi T, Ebrahimie E (2013) Genome-wide analysis of key salinity tolerance transporter (HKT1;5) in wheat and wild wheat relatives (A and D genomes). In Vitro Cell Dev Biol Plant 49: 97–106

Zeeshan M, Lu M, Sehar S, Holford P, Wu F (2020) Comparison of biochemical, anatomical, morphological, and physiological responses to salinity stress in wheat and barley genotypes deferring in salinity tolerance. Agronomy 10: 127

Zhang WD, Wang P, Bao Z, Ma Q, Duan LJ, Bao AK, Zhang JL, Wang SM (2017) SOS1, HKT1; 5 and NHX1 synergistically modulate Na^+^ homeostasis in the halophytic grass Puccinellia tenuiflora. Front Plant Sci 8: 576

Zhu M, Shabala S, Shabala L, Fan Y, Zhou MX (2016) Evaluating predictive values of various physiological indices for salinity stress tolerance in wheat. J Agron Crop Sci 202: 115– 124

